# EGFL7 promotes immune evasion in glioblastoma by interaction with integrin β_2_

**DOI:** 10.1101/2025.07.28.667281

**Authors:** Sukrit Mahajan, Philipp Abe, Fanny Ehret, Carina Fabian, Nora Heinig, Marc Gentzel, Rajinder Gupta, Julieta Aprea, Paul Warnke, Eirini Moysoglou, Beatrice Wasser, Stephan Grabbe, Andreas Dahl, Kathrin Barth, Marc Schmitz, Frauke Zipp, Ulrike Schumann, Mirko HH Schmidt

## Abstract

Glioblastoma is the most aggressive form of malignant brain cancer, characterized by an immunosuppressive microenvironment and immune evasion. Despite the success of immune checkpoint inhibitors in other cancers, immunotherapies such as anti-PD1 have shown limited efficacy in glioblastoma, underscoring the need to identify tumor-intrinsic mechanisms that sustain this immunosuppressive microenvironment and to develop more effective therapeutic strategies targeting them. Previously, the secreted factor epidermal growth factor-like protein 7 (EGFL7) has been shown to promote brain tumor growth by affecting the glioblastoma microenvironment (GME). However, its impact on the immune system remained enigmatic. Here, we studied the role of EGFL7 in shaping the immune landscape in glioblastoma and identified the underlying molecular mechanisms it engages to drive glioma immune evasion. Single-cell transcriptomic profiling of immune cells derived of glioblastoma revealed that EGFL7 promotes an immunosuppressive GME, characterized by enhanced T cell exhaustion and polarization of macrophages towards a protumorigenic state. Proteomic profiling of EGFL7’s interactome in glioma revealed its interaction with integrin β2 (ITGB2), an immune cell surface receptor involved in cell adhesion and migration. Mechanistic studies uncovered the central role of this interaction for immune evasion, which promoted T cell exhaustion and the polarization of macrophages towards a pro-tumorigenic state. Genetic perturbation of the EGFL7-ITGB2 axis attenuated immunosuppression and prolonged the survival of glioblastoma-bearing mice. Remarkably, a combinatorial regimen of anti-EGFL7 and the checkpoint inhibitor anti-PD1 improved the efficacy of this drug, which by itself did not improve glioma patient survival so far. In conclusion, our study provides unequivocal evidence that EGFL7 mediates immune evasion in glioma and has great potential to serve as an add-on drug target to improve immunotherapies not functional in glioblastoma patients so far.

## Introduction

Glioblastoma, the most aggressive form of primary brain tumors, is characterized by immune evasion, angiogenesis, and inevitable recurrence despite available treatment options for the primary tumor such as surgical resection plus radiotherapy and chemotherapy^1,2^. Consequently, the prognosis of affected patients remains dismal, with a median survival of 15-18 months post diagnosis upon standard-of-care treatment^2^. Previous studies have highlighted the critical role of immune evasion in glioblastoma. However, the molecular mechanisms enabling brain tumor cells to evade immune surveillance remain poorly understood.

The glioblastoma microenvironment (GME)^3,4^ is characterized by profound alterations to the healthy brain parenchyma and has been suggested as a major driver of immune evasion. It constitutes a hostile milieu promoting the exhaustion of T cells along with the polarization of macrophages towards a pro-tumorigenic fate leading to an anti-inflammatory state, which supports tumor growth, and prevents an efficient anti-tumor immune response^5–7^. To curb the impact of immunesuppressive GMEs and restore anti-tumor immune responses, immunotherapies involving adoptive T cell therapy, immune checkpoint inhibitors such as anti-PD1 or vaccines have been developed^8–11^. Despite demonstrating clinical efficacy in non-small cell lung cancer^12^, melanoma^13^ and other solid tumors^14^, anti-PD1 therapies have been largely ineffective in glioblastoma^15,16^, which makes it imperative to identify tumor-intrinsic factors and underlying mechanisms that maintain an immunosuppressive GME for developing more effective therapies.

In this context, epidermal growth factor-like protein 7 (EGFL7), a secreted proangiogenic factor^17,18^ deposited in the GME, has been described to influence CNS inflammation^19,20^. Mechanistically, EGFL7 downregulates key endothelial adhesion molecules such as ICAM1 and VCAM1, thus facilitating immune escape^21^. Moreover, EGFL7 contributes to glioma progression by promoting blood vessel formation and maturation within the tumor microenvironment^19^. While EGFL7 has been established as a modulator of the tumor vasculature in glioblastoma, its impact on immune regulation within the GME remains elusive.

In the current study, glioblastoma models were applied to investigate how EGFL7 shapes the immune landscape in malignant brain tumors. Single-cell RNA sequencing (scRNA-seq) has been employed in order to identify distinct immune cell populations affected by EGFL7. Subsequently, the localization of immune cells in glioblastoma specimens has been precisely mapped using spatial transcriptomics. Further, proteomic profiling of the intratumoral EGFL7 interactome by mass spectrometry, combined with mechanistic studies involving cell-based assays and mouse models, was leveraged to uncover molecular pathways engaged by EGFL7 to promote immune evasion in glioblastoma. These findings elucidate the role of EGFL7 in regulating glioma progression and open novel therapeutic avenues for glioblastoma treatment.

## Results

### Elevated EGFL7 levels correlate with decreased glioblastoma survival

To investigate the relevance of EGFL7 for glioblastoma patient survival, EGFL7 levels of IDH mutant (low-grade glioma) and IDH wild-type (high-grade glioma) specimens, collected in the Chinese glioma gene atlas (CGGA), were compared and unraveled a significant upregulation of EGFL7 in high grade glioma samples (Figure 1a; n = 213 vs 204, *p* = 3.1 x 10^-6^). Further, data derived of The Cancer Genome Atlas (TCGA) were correlated to EGFL7 expression using the R2: Genomics Analysis and Visualization Platform (http://r2.amc.nl). Interestingly, glioma patients with high EGFL7 levels displayed shorter survival compared to patients with low EGFL7 expression (Figure 1b; n = 68 vs 205 specimens, *p* = 0.074). Moreover, these patients displayed a rapid decline in survival during the latter half of the follow-up period. In conclusion, EGFL7 becomes upregulated in high grade glioma and increased EGFL7 levels correlate with poor clinical outcome in glioblastoma patients.

**Figure 1:**
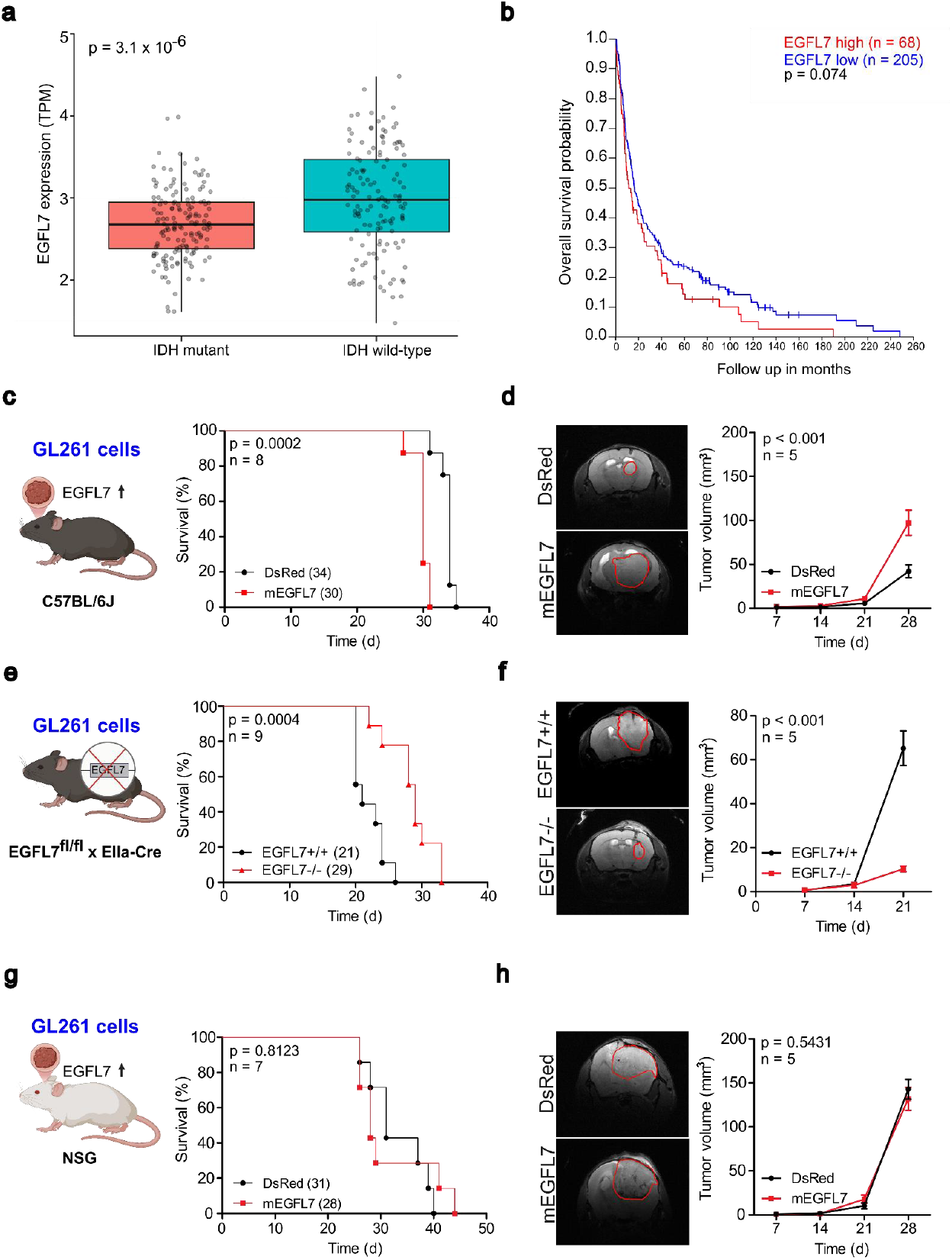
High levels of EGFL7 correlate with a poor prognosis in glioblastoma. **a:** Gene expression analysis using glioma patient samples derived of the Chinese glioma gene atlas (CGGA) reveals an increased expression of EGFL7 in IDH wild-type high-grade glioma compared to IDH mutant low-grade glioma. **b:** Kaplan-Meier curves of glioma patients from The Cancer Genome Atlas (TCGA) correlate high EGFL7 expression with reduced patient survival. **c:** Kaplan-Meier curves show decreased survival of mice bearing GL261-mEGFL7 gliomas compared to GL261-DsRed control tumors. **d:** Representative MRI scans 28 d post implantation (left). Longitudinal MRI analysis shows an increased tumor volume in GL261-mEGFL7 glioma-bearing mice compared to GL261-DsRed tumor-bearing control animals, 28 d post tumor implantation (right). **e:** Kaplan-Meier curves show prolonged survival of EGFL7-/- mice implanted with GL261-DsRed gliomas compared to tumor-bearing control littermates (right). **f:** Representative MRI scans 21 d post tumor implantation (left). Longitudinal MRI analysis shows decreased GL261-DsRed glioma volumes in EGFL7-/- mice compared to tumors grown in control littermates 21 d post implantation (right). **g:** Immunodeficient NSG mice bearing GL261-mEGFL7 gliomas or GL261-DsRed control tumors survive comparably long. **h:** Representative MRI scans 28 d post tumor implantation (left). Longitudinal MRI analysis in both groups shows comparable tumor sizes in GL261-mEGFL7 and GL261-DsRed glioma-implanted mice 28 d post tumor implantation (right). Statistical analyses were performed using log-rank test (**c**,⍰**e**,⍰**g**) and two-way ANOVA (**d**,⍰**f**,⍰**h**).

This correlation prompted further investigations using orthotopic experimental glioma models, in which the role of EGFL7 for glioma formation and survival was assessed using EGFL7 gain- and loss-of-function studies (Extended Data Figure 1a). In gain-of-function studies, GL261 glioma cells derived of C57BL/6J mice were engineered to ectopically express murine, V5-tagged EGFL7 (mEGFL7) and DsRed, both separated by a self-cleaving P2A peptide (Extended Data Figure 1b, top). GL261 cells solely expressing DsRed served as negative control. Upon retroviral infection, DsRed-expressing GL261 cells were collected by fluorescence-activated cell sorting (FACS) and the successful expression of V5-tagged mEGFL7 was verified by immunoblotting (Extended Data Figure 1b, bottom).

Subsequently, cells were intracranially implanted into the striatum of wild-type C57BL/6J mice and glioma growth was monitored until mice showed first symptoms. As a proof-of-principle for transgene expression, GL261-mEGFL7 and GL261-DsRed tumors were immunostained for V5-tagged mEGFL7, which was readily detectable upon ectopic expression (Extended Data Figure 1c). Remarkably, GL261-mEGFL7 glioma-bearing mice displayed a significant reduction in survival, with a median lifespan of 30 d compared to 34 d in GL261-DsRed control (Figure 1c; n = 8, *p* = 0.0002). Additionally, a significant increase in tumor volume was observed in GL261-mEGFL7 glioma-bearing mice compared to GL261-DsRed tumor-bearing controls (Figure 1d; 97 vs 42 mm^3^, n = 5, *p* < 0.001) 28 d post implantation as measured by MRI.

In the converse experiment, GL261-DsRed cells were intracranially implanted into the striatum of EGFL7 knock-out mice (EGFL7-/-) as well as control littermates (EGFL7+/+). The constitutive knock-out of EGFL7 prolonged the median survival of glioma-bearing mice from 28 d in EGFL7+/+ mice to 35 d in EGFL7-/- animals (Figure 1e; n = 10, *p* < 0.0001). Moreover, EGFL7-/- mice displayed comparatively smaller tumors in comparison to littermate controls (Figure 1f; 10 vs 65 mm^3^, n = 5, *p* < 0.001) 21 d post implantation.

To corroborate these findings in an alternative model, SB28 cells were subjected to comparable treatments. As a result, SB28-mEGFL7 glioma-bearing mice lived 5 d shorter compared to SB28-DsRed tumor-bearing control animals (Extended Data Figure 1d; 21 vs 26 d, n = 9, *p* = 0.0038). Moreover, SB28-mEGFL7 gliomas grew significantly bigger compared to SB28-DsRed tumors (Extended Data Figure 1e; 43.32 vs 13.46 mm^3^, n = 4, *p* = 0.0002) 21 d post tumor implantation. Conversely, EGFL7-/- mice lived 8 d longer compared to littermate controls upon SB28-DsRed implantation (Extended Data Figure 1f; 29 vs 21 d, n = 9, *p* = 0.0004) and displayed a reduced tumor volume (Extended Data Figure 1g; 10.54 vs 23.92 mm^3^; n = 5, *p* = 0.0052) 21 d post implantation.

In sum, EGFL7 promoted glioma growth in different experimental brain tumor models, which prompted an investigation of whether or not this phenotype had an immunological cause. Therefore, the gain-of-function model was applied to immunodeficient NSG mice. Remarkably, no significant difference in survival of mice implanted with GL261-mEGFL7 and GL261-DsRed cells was observed (Figure 1g; 28 vs 31 d, n = 7, *p* = 0.8123) and tumors grew to comparable sizes (Figure 1h; 143 vs 132 mm^3^, n = 5, *p* = 0.5439) 28 d post implantation as measured by MRI. Likewise, NSG mice implanted with SB28-mEGFL7 and SB28-DsRed glioma survived for comparable time frames (Extended Data Fig 1h; 27 vs 28 d, n = 6, *p* = 0.6047) and grew tumors of comparable size (Extended Data Figure 1i; 57 vs 56 mm^3^, n = 5, *p* = 0.9887) 28 d post tumor implantation.

In conclusion, EGFL7 affects glioma progression by modulation of the immune system.

### EGFL7 promotes immune evasion of glioma

In order to determine the cellular cause of EGFL7’s impact on the immune system, CD45^+^ immune cells were isolated by magnetic-activated cell sorting (MACS) from GL261-mEGFL7- and GL261-DsRed-derived glioma specimens 28 d post tumor implantation (Figure 2a, top). 42,289 cells were subjected to scRNA-seq analyses, resulting in 31 clusters from 8 samples, including 4 biological replicates from each group to account for tumor heterogeneity. Following quality control analysis, different clusters were predicted based on the nomenclature provided by García-Vicente et al.^22^ for clustering CD45^+^ immune cells into lymphoid and myeloid populations (Figure 2a, bottom). The lymphoid cluster was primarily composed of T cells, along with NK and a small amount of B cells, while the myeloid cluster displayed mostly macrophages, microglia and dendritic cells (Figure 2b).

**Figure 2:**
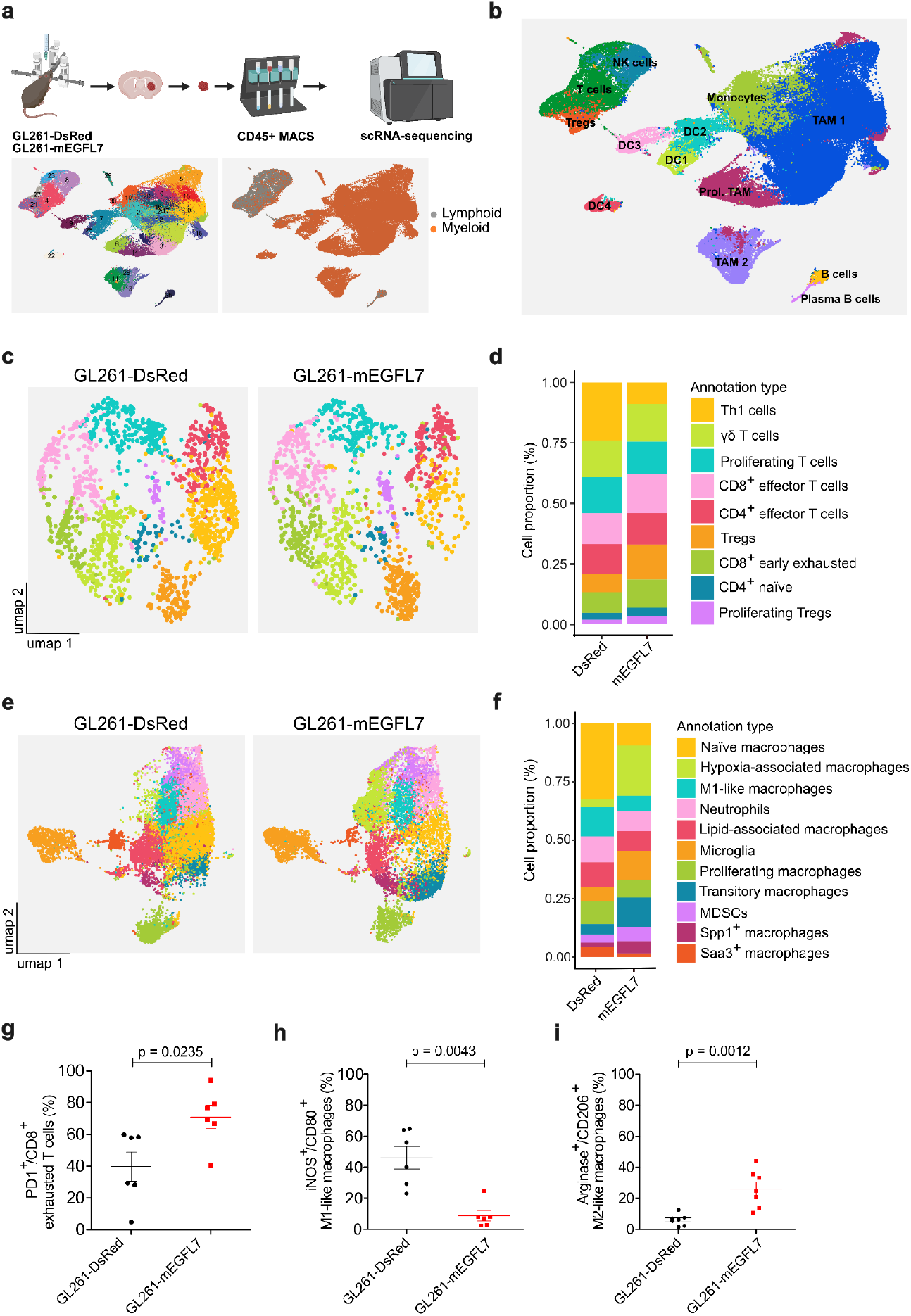
Ectopic EGFL7 promotes immune evasion of glioma. **a:** Schematic workflow of CD45^+^ immune cell isolation from GL261-mEGFL7 and GL261-DsRed-gliomas, followed by single-cell RNA sequencing (scRNA-seq) (top). In brief, CD45^+^ immune cells were isolated from tumor samples by MACS and analyzed using scRNA-seq, resulting in 31 immune cell clusters, which were classified into myeloid and lymphoid subsets based on markers defined by García-Vicente et al. (2025)^22^ (bottom). **b:** Clusters were further annotated on the basis of the immune cell atlas generated by Pombo Antunes et al. (2021)^23^. **c:** UMAPs of T cell clusters from GL261-DsRed and GL261-mEGFL7 gliomas. **d:** Stacked barplots show an increased proportion of CD8^+^ early exhausted T cells in GL261-mEGFL7 gliomas compared to GL261-DsRed tumors. **e:** UMAPs of myeloid clusters from GL261-mEGFL7 and GL261-DsRed gliomas. **f:** Stacked barplots showing a reduction in M1-like macrophages with concurrent increase in M2-like hypoxia-associated macrophages in GL261-mEGFL7 gliomas compared to GL261-DsRed tumors. Flow cytometry analyses verified that ectopic mEGFL7 induces g: more CD8^+^ exhausted T cells, **h:** less M1-like macrophages, and i: more M2-like macrophages in GL261-mEGFL7 gliomas compared to GL261-DsRed tumors (n ≥ 6). Statistical analysis was performed using the Mann-Whitney U test.

Different subtypes of T cells were identified in the T cell cluster based on known T cell markers (Figure 2c, Extended Data Figure 2a), such as CD8^+^ early exhausted T cells (*Pdcd1*^*+*^ *Lag3*^*+*^ *Gzma*^*low*^), CD8^+^ effector T cells (*Gzma*^*+*^ *Gzmb*^*+*^ *Gzmk*^*+*^) or CD4^+^ regulatory T cells (*Foxp3*^*+*^ *Il2ra*^*+*^). Proportional analysis of different subtypes revealed an increase in CD8^+^ exhausted T cells (11.7% vs 8.4%) in GL261-mEGFL7 glioma compared to GL261-DsRed controls (Figure 2d, Extended Data Figure 2b).

Analyses of the myeloid cluster according to the immune cell atlas for determining cell identity^23^ revealed macrophages, monocytes and microglia as the largest populations of myeloid residents in the glioma mass (Figure 2b+e, Extended Data Figure 2c). Remarkably, GL261-mEGFL7 tumors displayed a significant reduction in pro-inflammatory *CD80*^*+*^ *CD86*^*+*^ M1-like macrophages (6.75% vs 12.73%) compared to GL261-DsRed controls. Concurrently, a substantial accumulation of anti-inflammatory M2-like macrophages was observed in GL261-mEGFL7 tumors marked by *Arg1*^*+*^ *Mrc1*^*+*^ *Adam8*^*+*^ hypoxia-associated macrophages (21.5% vs 3.5%) and *SPP1*^*+*^ *Mrc1*^*+*^ macrophages (5.3% vs 1.66%) (Figure 2f, Extended Data Figure 2d).

In sum, mEGFL7 promotes an increase in pro-tumorigenic but a reduction in anti-tumorigenic immune cell populations in glioma.

In order to substantiate these scRNA-seq results, glioma resident immune cells were analyzed by flow cytometry, which verified an increase in *PD1*^*+*^ CD8^+^ exhausted T cells (Figure 2g; 71% vs 40%, n = 6, *p* = 0.0235) in GL261-mEGFL7 glioma specimens compared to GL261-DsRed controls. Additionally, a significant decrease in *CD80*^*+*^ *INOS2*^*+*^ M1-like pro-inflammatory macrophages (Figure 2h; 13% vs 40%, n = 7, *p* = 0.0043) was observed, alongside with a concurrent increase in pro-tumorigenic *CD206*^*+*^ *Arginase*^*+*^ M2-like macrophages (Figure 2i; 26% vs 6%, n = 7, *p* = 0.0012).

Likewise, SB28-mEGFL7 glioma tissue yielded a higher percentage of CD8^+^ exhausted T cells (Extended Data Figure 3a; 18% vs 6.9%, n = 8, *p* = 0.0004), a decrease in pro-inflammatory M1-like macrophages (Extended Data Figure 3b; 11% vs 31%, n = 8, *p* = 0.0006) and an increase in M2-like ones (Extended Data Figure 3c; 55% vs 36%, n = 9, *p* = 0.0054) compared to SB28-DsRed controls.

In conclusion, ectopic mEGFL7 promoted an anti-inflammatory and pro-tumorigenic immune environment in glioma.

### Loss-of-EGFL7 promotes anti-tumor immunity in glioma

Data above was corroborated by a loss-of-function study using a constitutive EGFL7 knock-out mouse model. GL261-DsRed cells were implanted into the striatum of EGFL7-/- animals or littermate controls and tumors were harvested after 28 d. Subsequently, CD45^+^ immune cells were enriched by MACS and were characterized by scRNA-seq as described above. Analysis of 46,411 cells resulted in 31 clusters from 8 samples, including 4 biological replicates from each group to account for tumor heterogeneity.

Most importantly, an approximate two-fold increase in *Gzma*^*+*^ *Gzmb*^*+*^ CD8^+^ effector T cells was found in glioma grown in EGFL7-/- mice (14.13% vs 7.2%) compared to littermate control tumors (Figure 3a, Extended Data Figure 4a), which is remarkable because these cells were found exhausted upon mEGFL7 expression as described above. Further, analysis of the macrophage/monocyte cluster revealed an increase in *CD80*^*+*^ *INOS2*^*+*^ M1-like macrophages (13.8% vs 11.9%) and a major decrease in immunosuppressive M2-like *Fabp5*^*+*^ *Gpnmb*^*+*^ *Lgals3*^*+*^ lipid-associated macrophages (10.67% vs 16.3%) in gliomas grown in the striatum of EGFL7-/- mice compared to littermate control specimens (Figure 3a+d, Extended Data Figure 4c+d).

**Figure 3:**
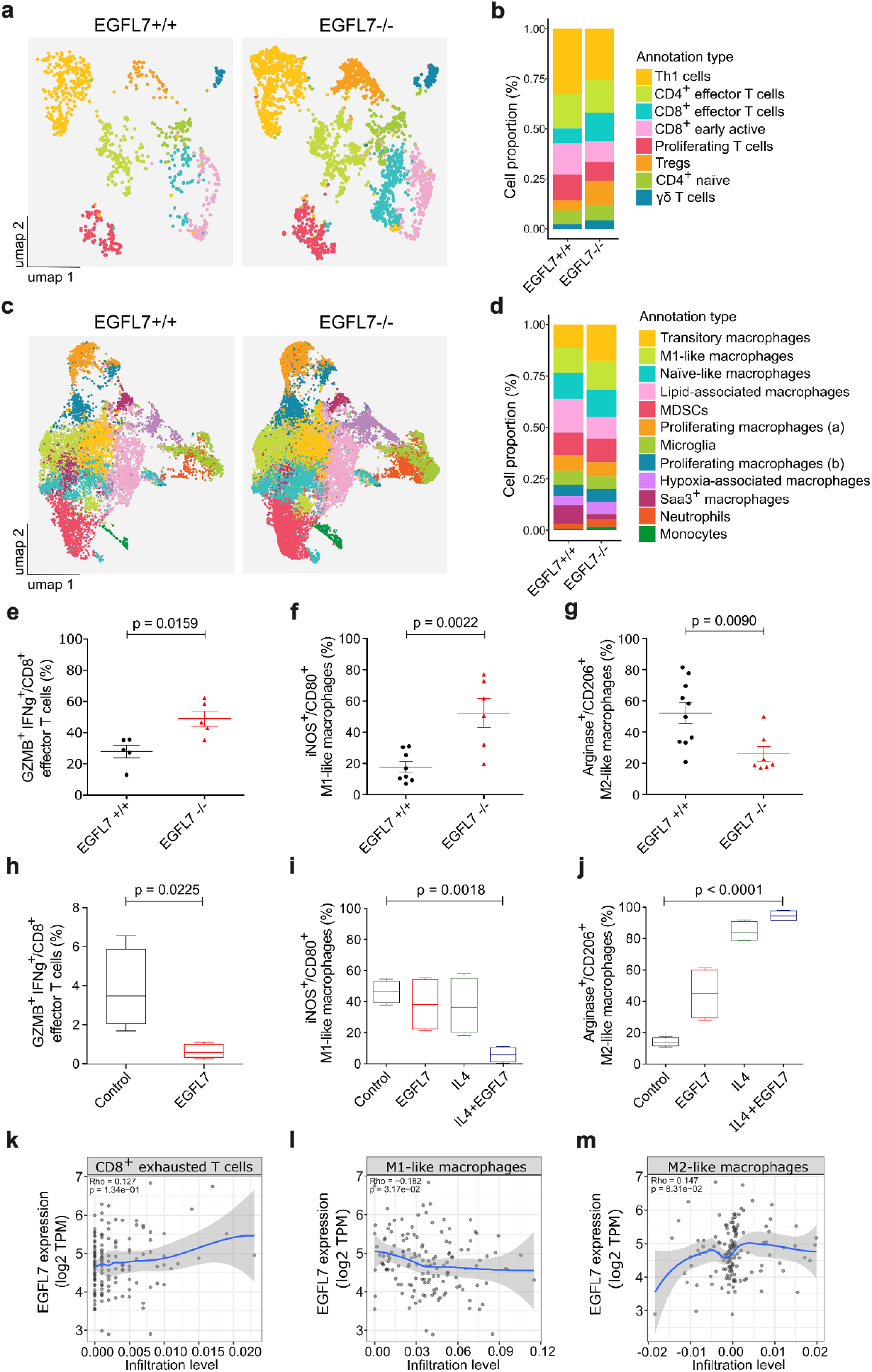
EGFL7 knock-out improves anti-tumor immunity in glioblastoma. **a-d:** UMAP visualizations and corresponding stacked barplots illustrate an altered distribution pattern of T cell (**a, b**) and myeloid cell (**c, d**) populations in GL261-DsRed glioma-bearing EGFL7-/- mice compared to tumors grown in control littermates. Subsequent flow cytometry analyses of gliomas revealed an increased amount of **e:** CD8^+^ effector T cells and **f:** M1-like macrophages, but **g:** fewer M2-like macrophages in tumors grown in EGFL7-/- mice compared to control littermates (n ≥ 5). **h:** T cells derived of spleens of healthy C57BL/6J mice were stimulated with anti-CD3 and anti-CD28 antibodies, before being challenged with purified recombinant mEGFL7. Flow cytometry analyses revealed a reduction of CD8^+^ effector T cells in the presence of mEGFL7 compared to control (n = 3). **i, j:** Macrophages were generated from bone-marrow derived monocytes of healthy C57BL/6J mice and were treated with purified recombinant mEGFL7, IL-4 or a combination of both to determine the impact on macrophage polarization compared to untreated controls. Flow cytometry analysis revealed **i:** a reduction of M1-like macrophages **j:** but an increase in M2-like macrophages in the presence of EGFL7 and IL4 (n = 4). **k, l, m:** In human glioblastoma specimens, analyses using TIMER revealed that EGFL7 expression correlates with the infiltration of (k) CD8^+^ exhausted T cells in a positive, (l) M1-like macrophages in a negative, and (**m**) M2-like macrophages in a positive manner.

In conclusion, regarding the immune landscape in glioma, scRNA-seq studies in EGFL7 knock-out mice draw an inversed picture compared to the mEGFL7 gain-of-function model above.

Proportions of immune cells in the GL261-DsRed tumor mass of EGFL7-/- mice and littermate controls were further quantified by flow cytometry. In particular, a significant increase in *Gzmb*^*+*^ *Infγ*^*+*^ CD8^+^ effector T cells was detected in EGFL7-/- mice compared to littermate controls (Figure 3e; 49% vs 28%, n = 5, *p* = 0.0159). In the myeloid population, more *CD80*^*+*^ *iNOS*^*+*^ M1-like (Figure 3f; 52% vs 18%, n = 7, *p* = 0.0022) but less *Arginase1*^*+*^ *CD206*^*+*^ M2-like macrophages (Figure 3g; 22% vs 55%, n = 7, *p* = 0.009) were detected within the GL261-DsRed tumor mass of EGFL7-/- mice compared to littermate controls.

Together, gain- and loss-of-function models both revealed that EGFL7 promotes an anti-inflammatory and pro-tumorigenic glioma microenvironment by inducing CD8^+^ T cell exhaustion and driving macrophage polarization from an M1-like towards an M2-like state. In order to show that this is a direct effect of EGFL7, T cells were isolated from spleens of healthy mice, and were stimulated using plate-bound anti-CD3 and anti-CD28 antibodies. Purified recombinant mEGFL7 was added to the culture medium and the percentage of CD8^+^ effector T cells was assessed after 5 d of incubation to measure T cell activation. In the presence of EGFL7, an almost 6-fold reduction was observed in *GzmB*^*+*^ *Ifnγ*^*+*^ CD8^+^ effector T cells compared to controls (Figure 3h; 0.64% vs 3.8%, n = 3, *p* = 0.0225), showing that EGFL7 directly promotes T cell exhaustion.

Likewise, the impact of EGFL7 on macrophage polarization was determined. Mouse hind leg bone marrow-derived macrophages (BMDMs) were incubated in a medium containing M-CSF along with purified recombinant mEGFL7 and interleukin-4 (IL4) to promote M2-like polarization. Cells were harvested after 24 h of incubation and flow cytometry analysis revealed that a combination of EGFL7 and IL4 caused an almost 8-fold reduction in M1-like *CD80*^*+*^ *iNOS*^*+*^ macrophages (Figure 3i; 5.75% vs 46.13%, n = 4, *p* = 0.0018). In parallel, the proportion of *Arginase1*^*+*^ *CD206*^*+*^ M2-like macrophages increased more than 3-fold in the presence of EGFL7 alone (Figure 3j; 45% vs 14%, n = 4, *p* < 0.0001) and about 6-fold when EGFL7 and IL4 were combined (Figure 3j; 94.68% vs 14%, n = 4, *p* < 0.0001).

In sum, EGFL7 creates an immune evasive environment in glioma by promoting T cell exhaustion and by shifting the polarization of macrophages away from an inflammatory M1-like towards a pro-tumorigenic M2-like state.

This prompted an investigation into whether EGFL7 affects these immune cell populations in human glioma as well. EGFL7 expression was correlated to the infiltration levels of T cells and macrophages in 287 glioblastoma patient samples obtained from TCGA, using the Tumor Immune Estimation Resource (TIMER)^24^. Analyses revealed a positive correlation of EGFL7 expression with exhausted T cell levels (Figure 3k; rho = 0.127, *p* = 0.134). Further, high EGFL7 expression levels negatively correlated with M1-like (Figure 3l; rho = -0.202, *p* = 0.018) but positively with M2-like (Figure 3m; rho = 0.246, *p* = 0.00377) macrophage infiltration.

In conclusion, findings above were translatable in the human setting with particularly strong correlations between EGFL7 levels and the infiltration of glioma by CD8^+^ exhausted T cells, M1- and M2-like macrophages.

### Spatial organization of immune cells in the glioma mass depends on EGFL7

In order to visualize the spatial localization of immune populations in the glioma mass, depending on EGFL7, GL261-DsRed cells were implanted into the striatum of EGFL7-/- mice and littermate controls. Brains were harvested after 25 d, cryo-sections were prepared and subjected to spatial transcriptomic analysis using 10x Visium. Spatial clusters in the brain sections were identified, and tumor tissue was further subdivided into tumor core (including tumor core and tumor shell 1) and tumor shell (comprising of tumor shell 2) to delineate the distribution of different immune cell populations. In general, immune cells were found throughout the glioma mass but were more enriched in the tumor shell and around the injection channel (Figure 4a).

**Figure 4:**
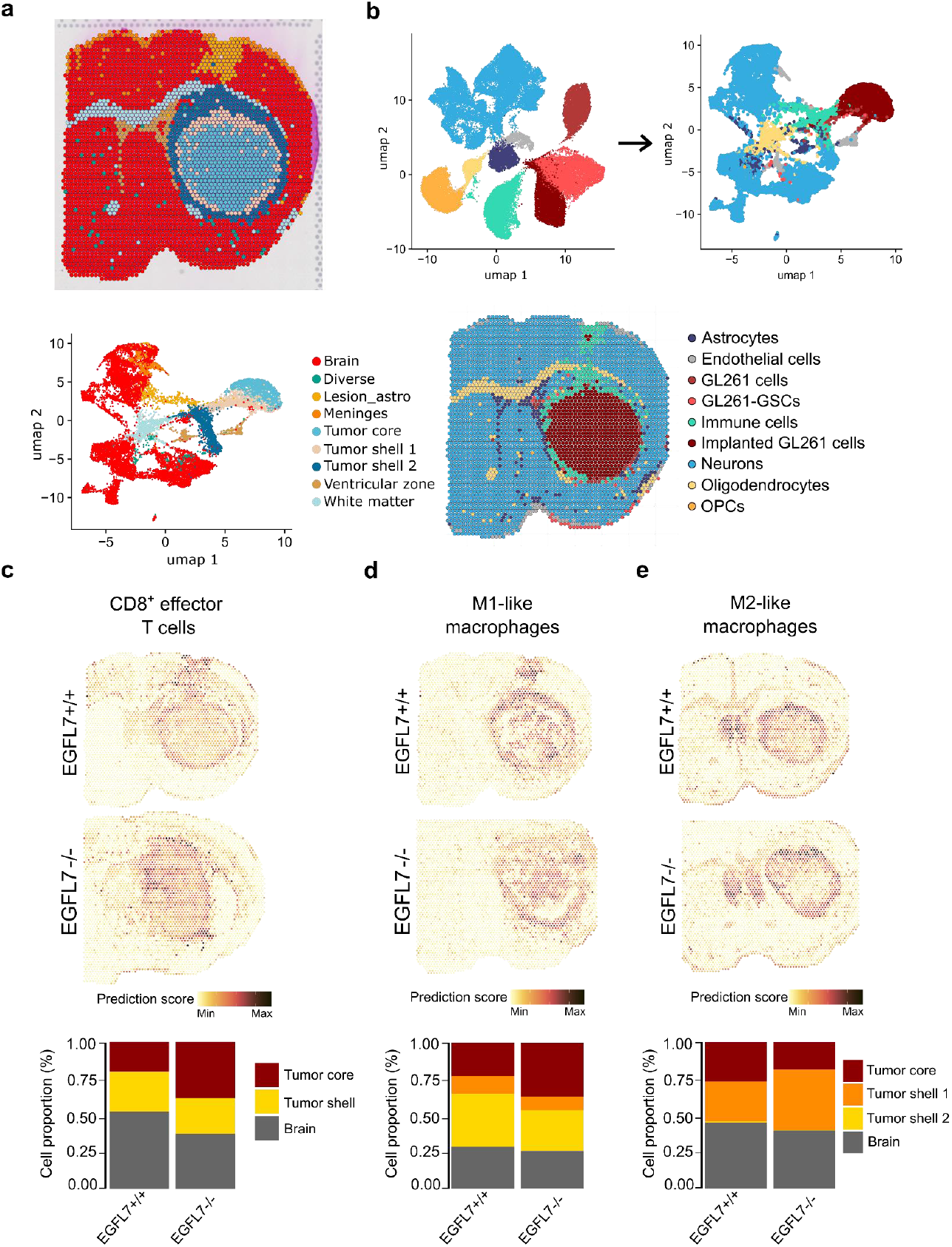
Spatial transcriptomics unravels the influence of EGFL7 on the distribution of immune cells in glioblastoma. **a:** Identification of spatial clusters in brain sections from GL261-DsRed gliomas grown in EGFL7-/- mice or littermate controls. **b:** Cell classification and spatial distribution of different cell types was based on annotations by Garcia Vicente et al. (2025)^22^. **c-e:** Distribution of immune cells in the glioma mass of EGFL7-/- mice or littermate controls using spatial transcriptomic analyses. Representative images (top) and quantitative analyses reveal **c:** an increased proportion of CD8^+^ effector T cells **d:** and M1-like macrophages but **e:** a decrease in M2-like macrophages in the mass of glioma implanted in EGFL7-/- mice compared to littermate control-derived tumor specimens (bottom).

Individual cell types such as immune cells, GL261 glioma cells, astrocytes and endothelial cells were predicted based on the study by García-Vicente et al.^22^ (Figure 4b). Prediction scores revealed a clear distinction among tumor and immune cell populations. T cells and macrophages exhibited distinct spatial heterogeneity across the tumor core, tumor shell, and neighboring healthy brain tissue. In particular, the distribution of T cells was affected by the absence of EGFL7 with a higher percentage of effector CD8^+^ T cells being present in the tumor core of EGFL7-/- mice compared to littermate controls (Figure 4c). Likewise, an elevation of M1-like macrophages was observed in the tumor core, suggesting that M1-like macrophages infiltrate the glioma mass more efficiently in the absence of EGFL7 (Figure 4d). Further, the tumor core displayed a much lower amount of pro-tumorigenic lipid-associated macrophages in EGFL7-/- mice compared to littermate controls, which were instead localized in tumor shell 1 (Figure 4e), showing that in the absence of EGFL7 less M2-like macrophages infiltrate the tumor mass.

In conclusion, spatial transcriptomics revealed that EGFL7 sequestered effector T cells and M1-like macrophages from the glioma mass while it caused an enrichment of immunosuppressive M2-like macrophages, such facilitating immune evasion and glioma progression.

### EGFL7 binds to ITGB2

In order to unravel how EGFL7 mediates immune evasion on the molecular level its interactome in glioma was determined upon immunoprecipitation of V5-tagged EGFL7 from GL621-mEGFL7 tumors. GL261-DsRed tumors served as negative controls due to the absence of a V5-tag, such accounting for any non-specific binding that may occur during the immunoprecipitation procedure. Proteins co-precipitating with EGFL7 were identified using liquid chromatography-based mass spectrometry in triplicate (Figure 5a). Particularly, EGFL7-binding proteins identified were involved in immune system processes as detected by Gene ontology analyses using GOrilla (Figure 5b). Among these proteins, various integrins were found including integrin α_4_ (ITGA4), integrin α_V_ (ITGAV), integrin β_1_ (ITGB1) and integrin β_2_ (ITGB2, also known as CD18) with ITGB2 being most abundant (Figure 5c) and known to be involved in immune cell adhesion and migration^25^.

**Figure 5:**
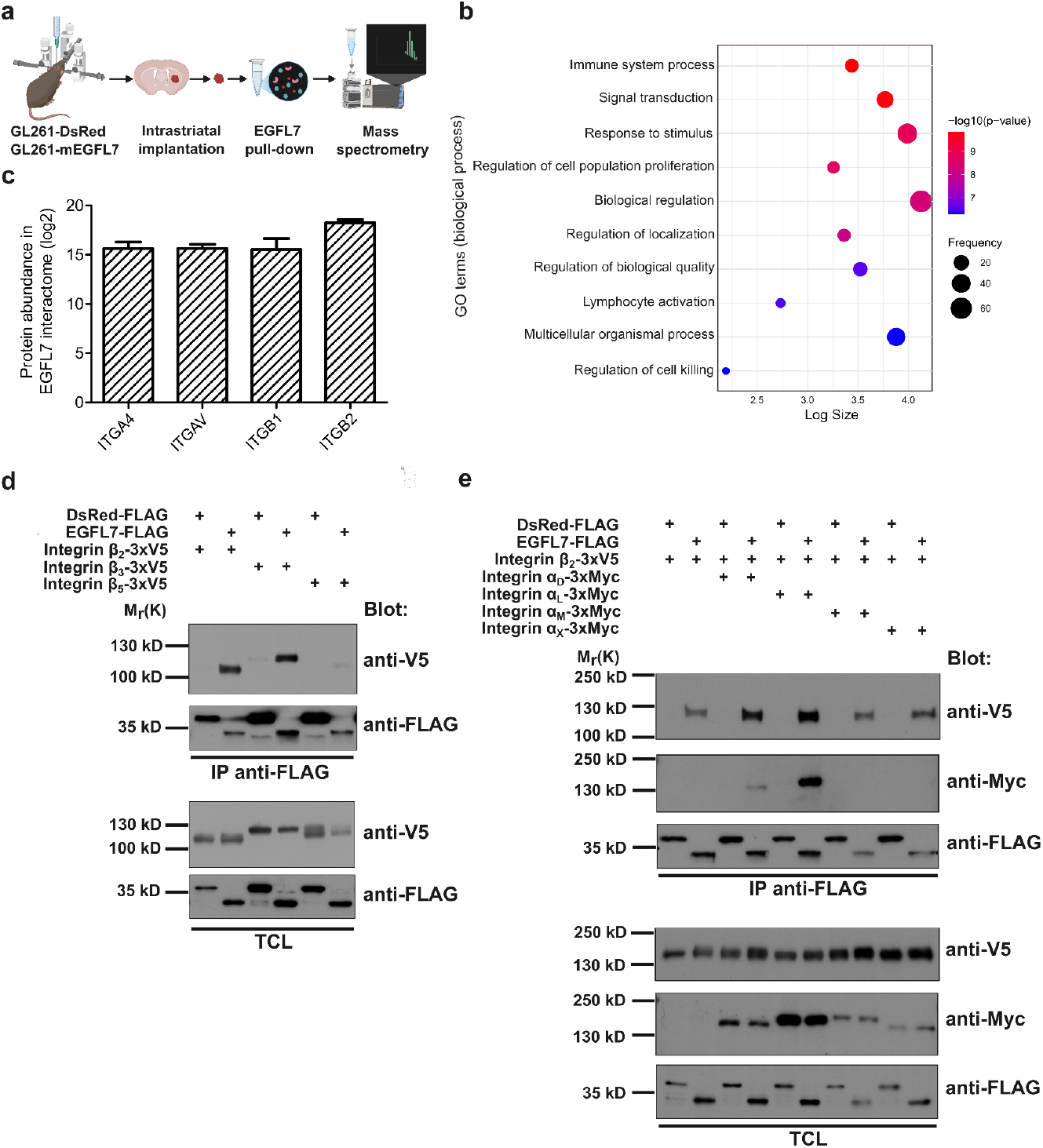
EGFL7 binds to ITGB2 in glioblastoma specimens. **a:** Schematic workflow of mass spectrometry analysis of the EGFL7 interactome in GL261-mEGFL7 and GL261-DsRed gliomas. V5-tagged EGFL7 was pulled down using an anti-V5 antibody (n = 3). **b:** Gene ontology analysis of proteins detected in the EGFL7 interactome. **c:** Barplot depicts the abundance of different integrins in the EGFL7 interactome. **d:** HEK293-EBNA cells were transfected with mammalian expression vectors encoding for Flag-tagged EGFL7 or DsRed as well as V5-tagged β-integrins. Cells were lysed and underwent co-immunoprecipitation analyses. Flag-tagged proteins were pulled down by anti-Flag. Immunoblots were performed using anti-Flag to detect DsRed and EGFL7 or anti-V5 to detect co-precipitating β-integrins. EGFL7 binds to ITGB2 as well as ITGB3, which served as a positive control. **e:** Plasmids encoding for Flag-tagged DsRed or EGFL7, V5-tagged β-integrins, and Myc-tagged α-integrins were transfected into HEK293-EBNA cells. Cells were lysed and underwent co-immunoprecipitation studies. Flag-tagged proteins were pulled down by anti-Flag antibody. Immunoblots were performed using anti-FLAG to detect DsRed and EGFL7, anti-V5 for β-integrins and anti-Myc for α-integrins. Data show that EGFL7 strongly associates with ITGAL-ITGB2, weakly with ITGAD-ITGB2 but barely with ITGAM-ITGB2 and ITGAX-ITGB2 complexes.

To confirm these interactions, putative binding partners were cloned into mammalian expression vectors. EGFL7 and DsRed were tagged with Flag, α-integrins with Myc and β-integrins with V5. Constructs were transfected into HEK293-EBNA cells, lysed and underwent co-immunoprecipitation studies using an anti-Flag antibody to pull down EGFL7 or DsRed as a non-binding control. The observed interaction of EGFL7 with ITGB2 and ITGB3 was about equally strong, while DsRed did not pull-down any of the integrins (Figure 5d). Previously, ITGB3 has been shown to bind to EGFL7^26^ and therefore served as a positive control. This prompted an investigation into which α-subunit may participate in the EGFL7-ITGB2 complex, because integrins exist as heterodimers comprised of α- and β-subunits. ITGB2 has been reported to dimerize with integrin-α_D_ (ITGAD), integrin-α_L_ (ITGAL), integrin-α_M_ (ITGAM) or integrin-α_X_ (ITGAX), which are also known as CD11a-d. Therefore, ITGB2 was co-expressed with each of these four α-subunits individually, along with EGFL7 or DsRed as a negative control. Co-immunoprecipitation studies using an anti-Flag antibody revealed a particularly strong interaction of EGFL7 with ITGAL-ITGB2, better known as lymphocyte function-associated antigen 1 (LFA-1), and a weaker one with ITGAD-ITGB2. However, a weak or no interaction was observed with ITAM-ITGB2 and ITAX-ITGB2 (Figure 5e). In sum, EGFL7 displayed a distinct interaction with ITGAL-ITGB2 highly expressed on T cells and involved in ICAM1-induced T cell activation.

### EGFL7 promotes glioma immune evasion in an ITGB2-dependent manner

In order to define whether EGFL7 affects immune evasion in glioma in an ITGB2-dependent manner, *ITGB2*^*fl/fl*^*;Scl-CreERT* mice were generated, allowing for the tissue-specific, tamoxifen-inducible knock-out of ITGB2 in hematopoietic stem cells (HSCs) and their immune cell progeny (Figure 6a). In brief, tamoxifen was administered to *ITGB2*^*fl/fl*^*;Scl-CreERT* mice or *ITGB2*^*fl/fl*^ litters for three consecutive days in order to generate ITGB2^ΔHSC^ mice and littermate controls. HSCs were allowed to generate immune cells for a period of eight wks before animals were sacrificed. Subsequently, CD45^+^ cells were isolated by MACS from the bone marrow of ITGB2^ΔHSC^ mice and littermate controls and were analyzed for *Itgb2* expression by quantitative reverse transcriptase-polymerase chain reaction (qRT-PCR). ITGB2^ΔHSC^ mice displayed a reduction in ITGB2 expression in CD45^+^ cells in the bone marrow by 94% compared to littermate controls (Figure 6b; n = 6, *p* = 0.0013). In CD45^+^ peripheral leukocytes, ITGB2 expression was reduced by 73 % (Figure 6c; n = 5, *p* = 0.0317), showing that the *Itgb2* locus got effectively recombined by Cre in the majority of HSCs and immune cells.

**Figure 6:**
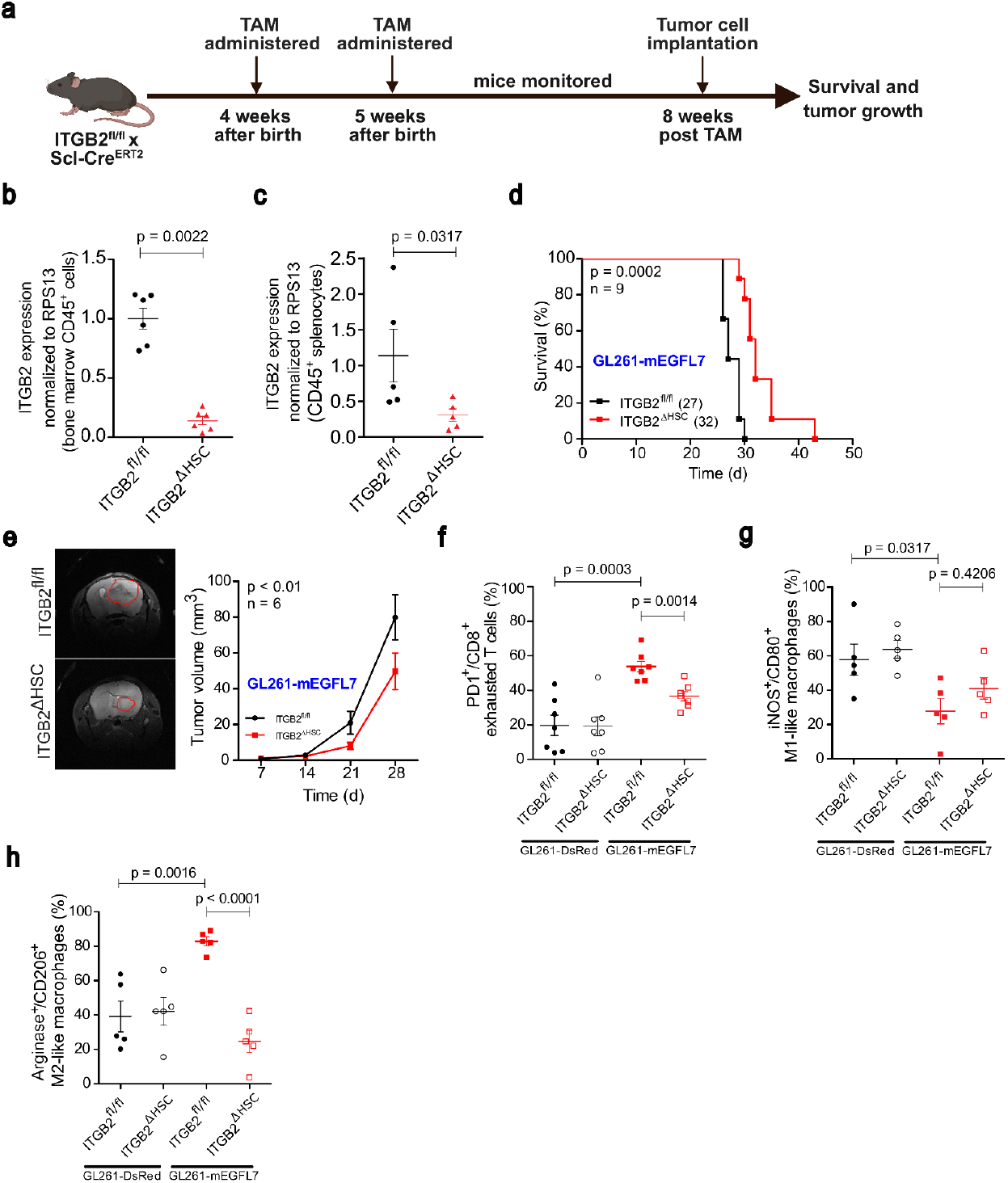
EGFL7 affects immune cells in an ITGB2-dependent manner. **a:** Schematic timeline showing tamoxifen administration at 4 and 5 wks of age to ITGB2^fl/fl^;Scl-CreERT mice to induce the deletion of ITGB2 in hematopoietic stem cells (HSCs), which later give rise to immune cells. 8 wks post tamoxifen treatment, an effective ITGB2 deletion was validated by qRT-PCR in **b:** bone marrow cells and **c:** splenocytes of ITGB2^ΔHSC^ mice compared to ITGB2^fl/fl^ littermate controls. **d:** 8 wks post tamoxifen treatment, GL261-mEGFL7 tumor cells were implanted in the striatum of mice. Kaplan-Meier curves show prolonged survival of glioma-bearing ITGB2^ΔHSC^ mice compared to tumor-bearing control littermates (log-rank test). **e:** Representative MRI scans, 28 d post GL261-mEGFL7 tumor implantation (left). Longitudinal MRI analysis revealed decreased volumes of tumors grown in ITGB2^ΔHSC^ mice compared to control littermates (two-way ANOVA; right). Flow cytometry analyses revealed an **f:** increased amount of CD8^+^ exhausted T cells, **g:** a reduction of M1-like and **h:** an increase in M2-like macrophages in GL261-mEGFL7 gliomas compared to GL261-DsRed tumors both grown in ITGB2^fl/fl^ control littermates. Knock-out of ITGB2 in ITGB2^ΔHSC^ mice rescued all these EGFL7 phenotypes.

Upon verification of the mouse model, GL261-mEGFL7 cells were intracranially implanted into the striatum of ITGB2^ΔHSC^ mice plus littermate controls and their survival was monitored until mice showed first symptoms. Most interestingly, ITGB2^ΔHSC^ mice survived 5 d longer compared to control littermates (Figure 6d; 32 vs 27 d, n = 9, *p* = 0.0002) and grew significantly smaller tumors (Figure 6e; 49.72 mm^3^ vs 79.88 mm^3^, n = 6, *p* < 0.01) 28 d post implantation as measured by MRI.

In conclusion, EGFL7 affects glioma growth depending on ITGB2 expressed on immune cells.

To define the mutual impact of EGFL7 and ITGB2 onto the immune landscape, GL261-mEGFL7 along with GL261-DsRed control cells were intracranially implanted into the striatum of ITGB2^ΔHSC^ mice and littermate controls. Mice were sacrificed after 28 d and CD45^+^ immune cells from the glioma masses were analyzed by flow cytometry. As expected, *PD1*^*+*^ CD8^+^ exhausted T cells were found increased (Figure 6f; 55% vs 20%, n =7, *p* = 0.0003), anti-tumorigenic *iNOS1*^*+*^ *CD80*^*+*^ M1-like macrophages decreased (Figure 6g; 27.8% vs 57.5%; *p* = 0.0317), and *Arginase*^*+*^ *CD206*^*+*^ M2-like macrophages increased (Figure 6h; 82% vs 39%, n = 5, *p* = 0.0016) in GL261-mEGFL7 tumors compared to GL261-DsRed specimens grown in littermate controls. However, each EGFL7 phenotype was rescued upon implantation of GL261-mEGFL7 cells in ITGB2^ΔHSC^ mice compared to specimens grown in littermate controls. As a result the proportions of *PD1*^*+*^ CD8^+^ exhausted T cells (Figure 6f; 33% vs 55%, n = 7, *p* = 0.0014), *iNOS1*^*+*^ *CD80*^*+*^ M1-like macrophages (Figure 6g; 41% to 27.8%; *p* = 0.4206) and *Arginase*^*+*^ *CD206*^*+*^ M2-like macrophages (Figure 6h; 24% vs 82%, n = 5, *p* < 0.0001) returned to levels comparable to control.

All in all, EGFL7 promotes glioma immune evasion in an ITGB2-dependent and an immune cell-specific manner.

### Anti-EGFL7 improves anti-PD1 glioma immunotherapy

Previously, anti-PD1 checkpoint blockade has been shown to reduce T cell exhaustion but yielded limited efficacy in glioblastoma^27^. Data above revealed that EGFL7 promotes T cell exhaustion, which prompted an investigation into whether blockage of EGFL7 alongside with anti-PD1 treatment might improve glioma immunotherapy.

Therefore, GL261-DsRed cells were implanted into the striatum of C57BL/6J mice and glioma growth was monitored until mice showed first symptoms. Mice received intraperitoneal injections of antibodies targeting EGFL7 (anti-EGFL7), PD1 (anti-PD1), a combination of both, or an isotype control (IgG) at d 16, 21, and 26 post tumor implantation (Figure 7a). As expected, anti-PD1 treatment did not prolong animal survival^28^. However, a combinatorial regimen combining anti-EGFL7 and anti-PD1 treatment improved the median survival of GL261-DsRed glioma-bearing mice up to 40 d compared to 33 d in IgG control (Figure 7b; n = 11, *p* = 0.0022). Further, a significant reduction in tumor volume was observed upon combinatorial treatment (Figure 7c; 27.24 mm^3^ vs 52.4 mm^3^, n = 6, p = 0.0222) 28 d post implantation as measured by MRI.

**Figure 7:**
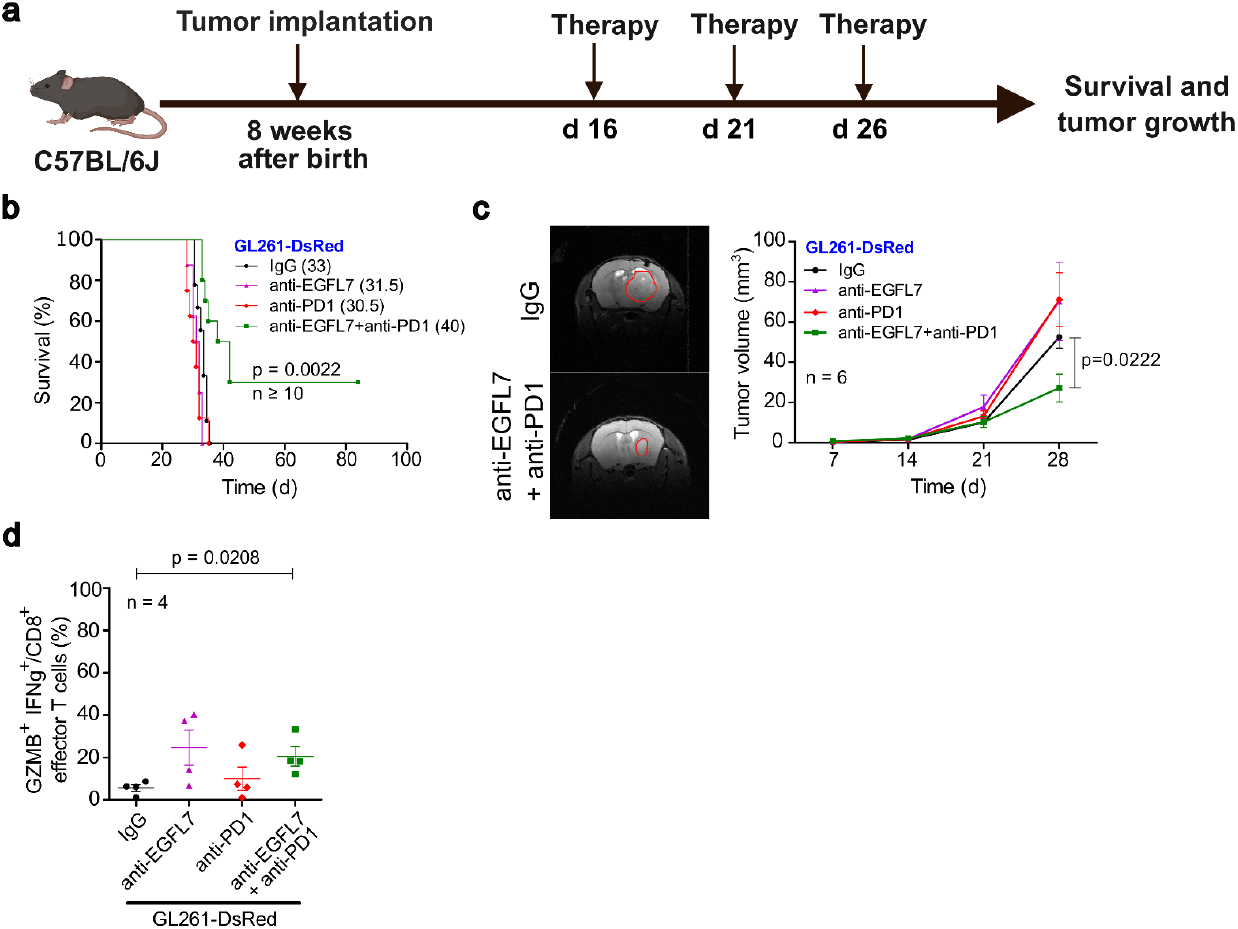
A combinatorial regimen of anti-EGFL7 and anti-PD1 improves glioma immunotherapy. **a:** Schematic describing the experimental paradigm in GL261-DsRed glioma-bearing mice treated with IgG control, anti-EGFL7, anti-PD1, or a combination of both at 16, 21 and 26 d post tumor implantation. **b:** Kaplan-Meier curves (n ≥ 8) show a significant survival benefit of animals treated with a combinatorial regimen of anti-EGFL7 and anti-PD1 compared to IgG (log-rank test). **c:** Representative MRI scans (left) and longitudinal MRI analysis 28 d post implantation (right) reveal reduced tumor volumes in GL261-DsRed implanted mice upon anti-EGFL7 and anti-PD1 combinatorial therapy compared to IgG controls (two-way ANOVA). **d:** CD8^+^ effector T cells were found increased about 4-fold in animals treated with a combinatorial regimen of anti-EGFL7 and anti-PD1 compared to IgG-treated controls as determined by flow cytometry analysis (Mann-Whitney U test).

Moreover, 28 d post tumor implantation, CD45^+^ immune cells from GL261-DsRed tumors were analyzed by flow cytometry. Quite remarkably, the combinatorial regimen of anti-EGFL7 and anti-PD1 led to a four-fold increase in *Gzmb*^*+*^ *Infγ*^*+*^ CD8^+^ effector T cells compared to IgG control treated mice (Figure 7d; 21% vs 5.6%, n = 4, *p* = 0.0208).

In conclusion, anti-EGFL7 may serve as a therapeutic strategy to improve anti-PD1 treatment-based immunotherapy of glioblastoma patients.

## Discussion

Glioblastoma remains incurable despite therapeutic advancements, with immune evasion playing a central role in tumor progression^29^. The tumor microenvironment constitutes an immunosuppressive milieu which plays an important role in evading the immune response. Tumor-associated macrophages (TAMs) in the GME release TGFβ and IL-10 to inhibit effector T cell function, thus promoting immunosuppression^30^. Further, TAMs express PD-L1 that binds PD1 on T cells and thus trigger T cell exhaustion^31^, which is favorable for glioma progression. In order to counteract these mechanisms of immune evasion, various immune checkpoint inhibitors have been developed. Unfortunately, their clinical efficacy in glioblastoma treatment remained ineffective^15,28,32^. In fact, anti-PD1 (nivolumab) therapy has been evaluated in clinical trials such as CheckMate 498 (NCT02617589)^33^ and CheckMate 143 (NCT02017717)^32^. However, it failed to improve overall survival in glioblastoma patients, which emphasizes the need to elucidate alternative mechanisms of immune evasion in glioblastoma.

Previously, we have shown that the neurovascular factor EGFL7, a protein largely secreted by endothelial cells^34^ but also neurons^35^, promotes glioma growth in a blood vessel-dependent manner^19^. Preliminary data obtained from brain tumor specimens used in this study offered the possibility that the immune landscape in glioma might be affected by EGFL7 as well. In particular, this seemed promising because we found that EGFL7 influenced CNS inflammation in a mouse model of multiple sclerosis termed experimental autoimmune encephalomyelitis^20^. Both studies offered the possibility that EGFL7 may act at the nexus of blood vessels and the immune system. As a matter of fact, one of the hallmarks of glioblastoma is extensive angiogenesis. Therefore, it seemed plausible that a protein secreted by endothelial cells and affecting the immune landscape might govern immune evasion mechanisms of malignant glioma, a hypothesis we challenged in this work.

Glioblastoma models *in vivo* revealed that ectopic EGFL7 markedly accelerated tumor growth and shortened the survival of immunocompetent mice. However, this pro-tumorigenic effect was not observed in immunodeficient mice, suggesting EGFL7 exerted its influence on glioma growth by engagement of a functional immune system. These observations align well with reports from other solid tumors, including breast and lung cancer, where EGFL7-mediated tumor growth was evident in immunocompetent but not immunodeficient mice^21^. This prompted an investigation into EGFL7’s impact on different immune cell populations across the glioma immune landscape.

Transcriptomic and functional analyses from gain- and loss-of-function models revealed that EGFL7 expression influenced the presence and activation state of several subsets of leukocytes, including T cells and macrophages. In particular, EGFL7 impaired effector T cell function and promoted their exhaustion. Concurrently, macrophages differentiated towards a pro-tumorigenic M2-like state at the expense of pro-inflammatory M1-like polarization in the presence of EGFL7. Spatial transcriptomics demonstrated an EGFL7-driven enrichment of M2-like macrophages in the tumor mass accompanied by restricted infiltration of CD8+ effector T cells and M1-like macrophages, data corroborated in patient-derived glioblastoma specimens. In this regard, accumulating evidence suggests that M2-like macrophages contribute to CD8^+^ T cell dysfunction and exhaustion in glioblastoma, thereby dampening effective anti-tumor immunity^36^. In addition, M2-like macrophages are known to transform the GME into an immunosuppressive state, by secreting immunosuppressive factors such as arginase and IL4^37^. These findings provide evidence that EGFL7 not only drives T cell exhaustion in a direct manner but also indirectly via M2-like polarization of macrophages while the M1-like state is suppressed. Collectively, this establishes a spatial and functional immunosuppressive environment that supports immune evasion and glioblastoma progression.

However, the molecular mechanisms driving EGFL7’s impact on T cells and macrophages remained enigmatic. To elucidate them, EGFL7’s protein interactome in glioma specimens was mapped by mass spectrometry. Most interestingly, ITGB2, an integrin expressed on the surface of immune cells, was identified as a primary interaction partner. EGFL7 showed preferential binding of ITGAL-ITGB2, an integrin heterodimer highly expressed on T cells^38^, while its interaction with ITGAD-ITGB2, mostly expressed on macrophages^39^, was much weaker. In brief, ITGB2 governs immune cell functions such as T cell activation, migration and cytokine secretion, along with macrophage polarization^38,40^. Thus, EGFL7 with its strong preference for ITGAL-ITGB2 inhibits the infiltration of effector T cells into the glioma mass, as observed in our spatial transcriptomics analyses.

EGFL7 may directly block the interaction of ITGAL-ITGB2 with ICAM1 by preventing an activating conformational change in ITGAL-ITGB2, as described for GDF15 in melanoma^41^. Similarly, EGFL7 may abolish the interaction between ITGAD-ITGB2 and its ligand VCAM1, thus restricting the infiltration of M1-like macrophages into the glioma mass, which express high levels of ITGAD-ITGB2 in contrast to M2-like macrophages^42,43^. Genetic ablation of ITGB2 in immune cells abolished EGFL7-induced T cell exhaustion, supporting a mechanistic link between EGFL7 and ITGB2-driven immunosuppression. Therefore, in addition to limiting immune cell infiltration, EGFL7 likely sustains immunosuppression through ITGB2-mediated downstream signaling. In this context, specific pathways downstream of ITGB2, such as YAP/TAZ signaling, may be activated, which has previously been shown to impair T cell function and to enhance PD-L1 levels in breast and lung cancer cells^44^. Furthermore, ITGB2 inhibits NFκB activation and promotes immune tolerance in macrophage populations^45^. This mechanism might very well be engaged by EGFL7 to prevent the polarization of macrophages towards an M1-like state, as it has been shown to downregulate NFκB signaling^43^. Plus, EGFL7 activates STAT3^46^, a key signal in M2-like macrophage polarization^47^ which could promote pro-tumorigenic differentiation.

Based on our findings, we propose a model wherein EGFL7 promotes immune evasion by its interaction with ITGB2. We suggest that during the early phase of glioblastoma development, different immune cell populations infiltrate the glioma mass, including T cells and macrophages. EGFL7 secreted into the GME by endothelial cells prevents the entry of T cells while also promoting T cell exhaustion via ITGAL-ITGB2. In addition, EGFL7 polarizes M1-like macrophages towards an M2-like state, through its interaction with ITGAD-ITGB2. M2-like macrophages are known to exacerbate T cell exhaustion by secreting arginase, which depletes arginine in the tumor microenvironment and thus suppresses T cell proliferation and cytokine production^48^. Moreover, since M2-like macrophages promote angiogenesis^49^, this will result in a positive feedback loop, causing elevated levels of EGFL7 in the GME, thereby further promoting M2 polarization as well as T cell exhaustion. Eventually, high levels of EGFL7 will result in maximal immune evasion and blood vessel formation, thereby driving aggressive tumor growth of glioblastoma (Figure 8).

**Figure 8:**
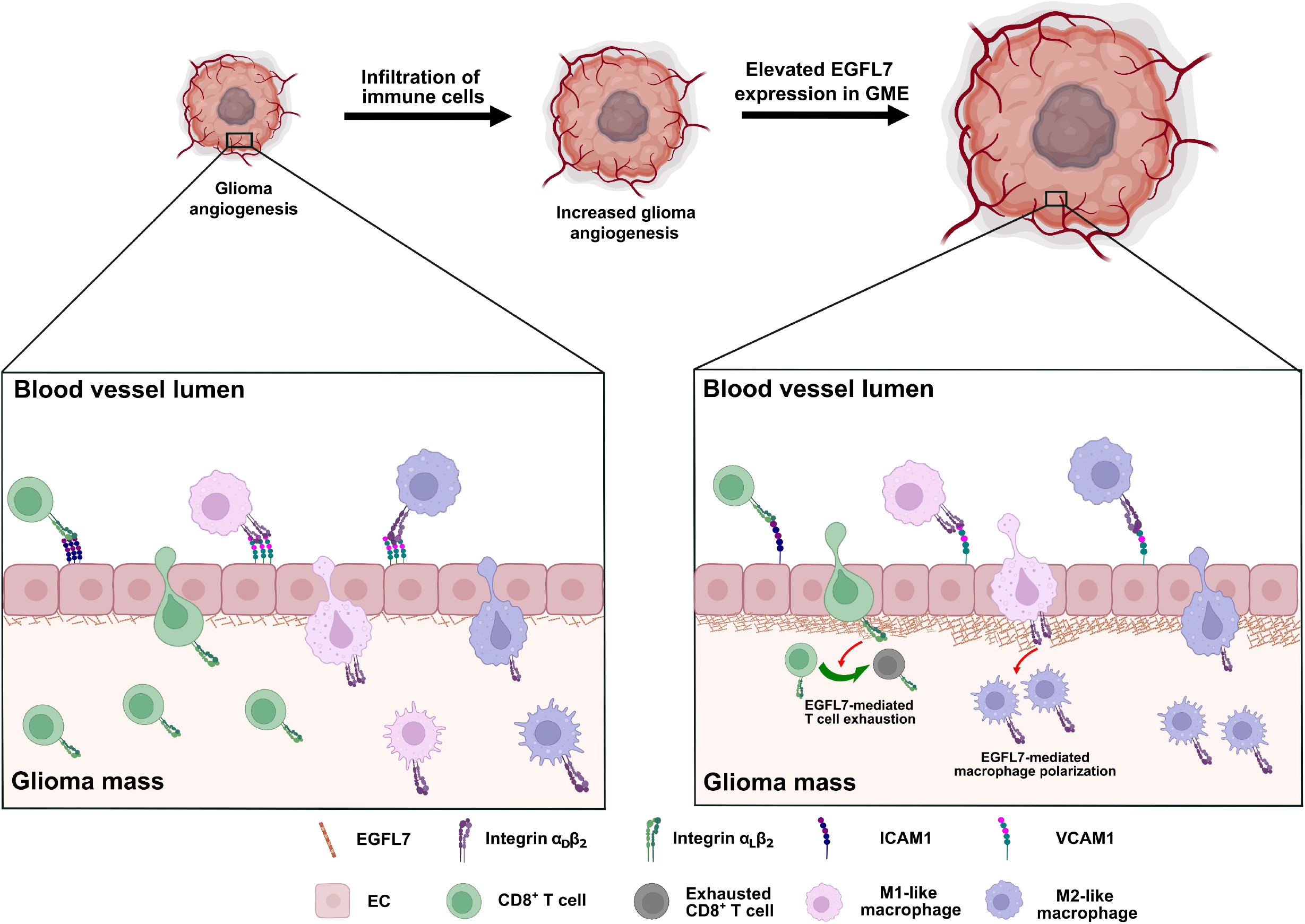
Schematic of how EGFL7 promotes immune evasion in glioblastoma. We suggest the following model. EGFL7 is secreted by endothelial cells in glioma angiogenesis. At low EGFL7 levels, the infiltration of immune cells into the glioma mass remains mostly unaffected. Once glioma proliferation accelerates, angiogenesis increases and local EGFL7 levels rise, which results in a downregulation of ICAM1 and VCAM1 expression on endothelial cells. EGFL7 binds ITGAL-ITGB2 on T cells and ITGAD-ITGB2 on M1-like macrophages, thereby reducing ICAM1 and VCAM1-binding, respectively. As a consequence, the infiltration of the glioma mass by these immune cells gets reduced. M2-like macrophages are less effected since they express lower levels of ITGAD-ITGB2. Further, EGFL7 promotes T cell exhaustion and polarizes macrophages infiltrating the glioma mass towards a pro-tumorigenic M2-like state by its interaction with ITGB2. Thus, T cells and M1-like macrophages are stalled at the tumor interface, while M2-like macrophages colonize the tumor mass. This way, the EGFL7-ITGB2 interaction promotes immune evasion and enhances glioma progression.

Thus, targeting the EGFL7-ITGB2 axis may prove beneficial in alleviating immune evasion in glioblastoma. However, the direct inhibition of ITGB2 to disrupt the EGFL7-ITGB2 axis might cause severe side effects since ITGB2 is known to regulate T cell activation along with functions of various other immune cells^50^. To bypass this shortcoming, we combined EGFL7 inhibition by parsatuzumab^51^ with anti-PD1 therapy to enhance the treatment efficacy of this particular immunotherapy. Data revealed that a combinatorial regimen of anti-EGFL7 and anti-PD1 not only improved glioma survival but also enhanced the anti-tumor immune response of glioma-bearing mice. Therefore, dual blockade of EGFL7 and PD1 overcame current limitations of PD1-based glioma immunotherapies.

In conclusion, our study identifies EGFL7 as a major driver of immune evasion in glioma, which promotes T cell exhaustion and macrophage polarization away from an inflammatory M1-like towards a pro-tumorigenic M2-like state by engaging ITGB2 signaling. Knock-out of ITGB2 in immune cells abolished this phenotype and blocking EGFL7 alongside with an immune checkpoint inhibitor improved the efficacy of anti-PD1 immunotherapy. These findings open a new and desperately needed avenue for improving the therapeutic outcome of glioblastoma patient treatment.

## Acknowledgements

We thank Nathalie Franke, Nicole Feind, Ella Herberger, Maria Lamm, Janet Lips, Stephanie Meyer, Anja Neißer, Martina Pinkert, and Doreen Streichert for excellent technical assistance; Susanne Weiche and Ellen Geibelt (CMCB, Dresden, Germany) for their support; and Tatyana Grinenko (Institute of Molecular Medicine, Shanghai, China) and Kurt Reifenberg (DKFZ, Heidelberg, Germany) for sharing tools.

This research was funded by the Deutsche Forschungsgemeinschaft (DFG, German Research Foundation) – Project-ID 318346496 – SFB 1292, project TP09 (FZ and MHHS) and the DFG grant SCHM2159/7-1 (MHHS).

## Author contributions

SM, PA, and FE designed and performed experiments; SM, PA, FE, CF, EM and US analyzed data; NH, KB, MG, RG, JA, PW, BW, SG, AD and MS contributed expertise and tools; FZ, US and MHHS designed and supervised the study and edited the manuscript.

## Conflict of interest

The authors declare that there are no conflicts of interest.

## Materials and Methods

### Animal models

Animal experiments were approved by the ethics committee of TU Dresden and conducted according to the German Animal Welfare Act. Experimental procedures were authorized by the Dresden Regional Council (Approval No. 25-5131/522/32 and 25-5131/542/35). All mice were housed under pathogen-free conditions with a 12 h light-dark cycle and free access to food and water at the Experimental Center (Medical Theoretical Center, Faculty of Medicine, Dresden, Germany).

C57BL/6J wild-type mice were purchased from Janvier Labs (Le Genest-Saint-Isle, France) and immunocompromised NOD/SCID/IL2rγ^null^ (NSG) mice originated from the Jackson Laboratory (Bar Harbor, ME, USA; stock no. 005557). Constitutive EGFL7 knock-out mice (*EGFL7*^*fl/fl*^*;EIIa-Cre*) were generated as previously described^54^. Briefly, EGFL7^fl/fl^ animals were crossed with EIIa-Cre pan-deleter animals expressing Cre under the adenovirus promoter EIIa^52^, allowing for the deletion of the EGFL7 exons 3 to 7 in germ line cells to create EGFL7-/- mice and EGFL7+/+ littermate controls.

Further, a conditional and tissue-specific ITGB2 knock-out mouse model was generated, enabling tamoxifen-inducible deletion of *Itgb2* from HSCs and their immune cell progeny. ITGB2^fl/fl^ mice (exon 3 of *Itgb2* flanked by loxP sites)^53^ were crossed with Scl-CreERT mice^54^ (*ITGB2*^*fl/fl*^*;Scl-CreERT*). At 4 and 5 wks of age, 150 mg per kg body weight (mpk) tamoxifen was administered i.p. for three consecutive days to induce the deletion of ITGB2 in HSCs (ITGB2^ΔHSC^), which give rise to immune cells. Tamoxifen-treated ITGB2^fl/fl^ mice served as littermate controls. A period of 8 wks was given for the repopulation of secondary lymphoid organs by mature ITGB2-deficient immune cells.

### Orthotopic glioma models

For stereotactic intracranial tumor injections, 8-10 wks old mice were anesthetized using 2% isoflurane (Primal) and were fixed in a stereotactic frame under constant anesthesia. Carprofen was administered as an analgesic at 10 mpk subcutaneously before surgery. A hole was drilled into the skull at the following coordinates relative to the bregma: 0.5 mm anterio-posterior and 2 mm medio-lateral. 25,000 GL261 or 1,000 SB28 cells were injected 3.5 mm dorso-ventral of the dura mater into the striatum of different mouse strains. Mice were euthanized by i.p. administration of an overdose of ketamine-xylazine (ketamine 450 mpk, xylazine 48 mpk), followed by transcardial perfusion with HBSS after 28 d or upon the development of glioblastoma-specific symptoms, such as weight loss, difficulty in movement and disheveled fur. The orthotopic tumor volume was measured using a 7 Tesla ClinScan 70/30 small animal ultra-high-field magnetic resonance imaging (MRI) scanner equipped with a mouse whole-body coil (Bruker, Ettlingen, Germany). T2-weighted images were acquired weekly (repetition time 2,500 ms, echo time 33 ms, slice thickness 0.5 mm) and were analyzed using ImageJ software.

For immunotherapy, isotype control (IgG2b), anti-EGFL7 (both from Genentech, San Francisco, USA), anti-PD1 (BioXcell, Hessen, Germany), or a combination of the latter were administered i.p. at a dose of 10 mpk to glioma-bearing mice on day 16, 21, and 26 post tumor implantation.

### Quantitative reverse transcriptase-polymerase chain reaction (qRT-PCR) analyses

RNA was isolated using the RNeasy PLUS Mini Kit (Qiagen). cDNA was synthesized with the RevertAid H Minus First Strand cDNA Synthesis Kit (Thermo Fisher Scientific) using oligo dT primer. qRT-PCR was performed using SsoAdvanced™ Universal Inhibitor-Tolerant SYBR Green Supermix (Bio-Rad Laboratories) on a CFX96 Touch Real-Time PCR Detection System (Bio-Rad Laboratories GmbH). Gene expression of ITGB2 was normalized to house-keeping gene ribosomal protein S13 (RPS13).

### Cell culture and purification of recombinant EGFL7

Mouse glioma cells GL261 and SB28 were cultured in Dulbecco’s modified Eagle’s medium (DMEM) supplemented with 10% fetal calf serum plus penicillin/streptomycin. HEK293-EBNA cells were cultured in DMEM supplemented with 10% fetal calf serum plus geneticin (all materials from Fisher Scientific, Waltham, MA, USA). Cells were cultured at 37 °C and 5% CO_2_. Recombinant mouse EGFL7 protein was expressed in Sf9 cells using a baculovirus system as described previously^55^.

### Viral transductions

Retroviral particles were produced by co-transfecting HEK293-EBNA cells with the vector RV-C207-CAG-dsRedExpress2-WPRE or RV-C207-dsRed-mEGFL7, together with the packaging plasmid pUMVC and the envelope plasmid pCMV-VSV-G, as previously described^56^. The resulting viral particle-containing supernatant was used to infect GL261 or SB28 cells. Successful gene transduction was visualized by DsRed expression and DsRed-positive cells were isolated by cell sorting using a FACS Aria II device (Becton Dickinson, Heidelberg, Germany).

### Immunoprecipitation studies and western blotting

For immunoprecipitation, cells were lysed in ice-cold lysis buffer (50 mM HEPES (pH 7.4), 150 mM NaCl, 10% glycerol, 1.5 mM MgCl_2_, 1% Triton X-100, 1 mM EDTA, 1 mM EGTA, 1 mM sodium orthovanadate, 20 mM NaF, 10 μM ZnCl_2_). Lysates were cleared at 4 °C by centrifugation at 16,000 g for 20 min. Subsequently, lysates were incubated with antibodies for 4 h at 4 °C under constant rotation. Protein-antibody complexes were collected by incubation with protein G magnetic beads (Invitrogen) for another 2 h, followed by centrifugation and washing the samples with wash buffer (lysis buffer without 1% Triton X-100). Samples were subsequently analyzed by mass spectrometry or processed further for western blotting. For western blot analysis, immunoprecipitates were boiled at 95 °C for 5 min in Laemmli buffer. Samples were loaded on an SDS-polyacrylamide (12%) gel, proteins were resolved by SDS-PAGE and transferred onto activated PVDF membranes. Anti-Flag (Proteintech), anti-V5 (Abcam), anti-Myc (Abcam) and anti-β-actin (Li-Cor) primary antibodies were used for immunodetection. Fluorescently conjugated IRDye 800CW donkey-anti-rabbit and IRDye 680RD goat-anti-mouse antibodies were used as secondary antibodies and detected using the LI-COR Odyssey device and LI-COR software (all from LI-COR Biosciences, Lincoln, NV, USA).

### Mass spectrometry analyses

Co-immunoprecipitated proteins bound to Protein G magnetic beads (Invitrogen) were processed for mass spectrometry analysis. In brief, protein-bound magnetic beads were resuspended in 2x-S-TRAP buffer (100mM TEAB, 10% SDS) and 1x wash buffer (50 mM Tris-HCl, 50 mM Tris-HCl, 150 mM NaCl, 1 mM EDTA, 10% v/v Glycerol). Eluted proteins were reduced, alkylated and digested according to the manufacturer’s protocol for S-Trap columns (Protifi, Fairport NY, USA). For analysis, samples were injected into the UHPLC system coupled to a Q-Exactive HF mass spectrometer (ThermoScientific, Bremen, Germany) and loaded onto a trap column in 0.1% formic acid (solvent A) at a flow rate of 3 µl per min. Subsequently, the separation column operated at a flow rate of 200 nl per min. Following equilibration with solvent A, peptides were separated in a linear gradient from 0% to 60% solvent B (60% acetonitrile, 0.1% formic acid). The mass spectrometer was operated in data-independent (DIA) acquisition mode. Peptide and protein identification was performed with DIA-NN V1.8 ^57^, using the mouse protein database. mEGFL7-interacting proteins detected in mass spectrometry were analyzed for Gene Ontology (GO) enrichment using the GOrilla tool^58^. The protein list was uploaded in “single ranked list” mode (*Mus musculus* background). GO biological processes were assessed with default settings (FDR q-value < 0.05). Enriched terms were visualized as a dot plot with colour intensity proportional to -log10 (p-value).

### Flow cytometry analyses

Glioma-bearing mice were euthanized, perfused and the tumor tissue processed as described above. Upon dissociation, cells were disseminated using a 70 µm strainer. Single cell suspensions underwent debris removal (Miltenyi Biotec), were washed and stained by Zombie aqua fixable live-dead dye (BioLegend). T cells were identified using the following markers: CD45-BUV805, CD3-BUV563, CD8-BUV395, Ly6C-BV605, CD279-BUV615, IFNg-BV711, GranzymeB-BB755-P, Foxp3-APC (BD Biosciences); CD4-FITC, CD25-PE-Cy7 (BioLegend). Myeloid cells were identified using the following markers: CD45-BUV805, CD11b-BUV395, CD80-BV421 (BD Biosciences); CD206-BV650, F4/80-BV786, iNOS-BB755-P (BioLegend); TMEM119-PE-Cy7, Arginase-PE (Invitrogen). Upon extracellular staining, cells were fixed and permeabilized before staining for intracellular markers, were recorded using a Symphony A3 device (Becton Dickinson) and were analyzed with FlowJo software (BD Biosciences).

### T cell stimulation assay

C57BL/6J mice were euthanized and perfused as described above. Spleens were dissociated according to the gentleMACS™ protocol “Neural Tissue Dissociation Kit” provided by Miltenyi, using an Octo MACS dissociator. For sorting T cells, the Pan T cell isolation kit II was used according to the manufacturer’s instructions (Miltenyi Biotec). 150,000 T cells per well were stimulated for 4 h at 37 °C using anti-CD3- and anti-CD28-coated 96-well plates. Cells were incubated with or without 1 µg/ml recombinant mEGFL7 protein, which was purified as described previously ^55^. After 5 d of incubation at 37 °C, cells were stained for different T cell markers and were analyzed using a Symphony A3 device (BD Biosciences).

### Macrophage polarization assay

C57BL/6J mice were euthanized and femur, tibia plus pelvis were taken for isolating bone marrow-derived macrophages (BMDMs). Cells were centrifuged, incubated in red blood lysis buffer, washed with PBS, and incubated at 37 °C in BMDM medium (RPMI 1640 supplemented with 10% FCS, 1% penicillin/streptomycin, 1% sodium pyruvate, 25⍰mM HEPES buffer and 2 mM L-glutamine) plus 50 ng/ml M-CSF (Invitrogen) for 10 d. 1 million cells per ml were cultured in the presence or absence of EGFL7 (1 μg/ml) and/or IL4 (20 ng/ml) for 24 h at 37 °C. Cells were harvested, stained for myeloid markers (see above) and analyzed by flow cytometry using a Symphony A3 device (BD Biosciences).

### Isolation of immune cells for scRNA-seq

Glioma-bearing mice were euthanized and perfused as described above. Tumors were excised and added to an enzyme mix (2.5 mg/ml collagenase D, 11 µg/ml pancreatic DNase, 0.5 µg/ml Trypsin inhibitor, 27 µg/ml Anisomycin, 50 mM HEPES) followed by dissociation using an Octo MACS dissociator and the NTDK protocol for 22 min at 37⍰°C. The reaction was stopped with ice cold HBSS followed by passing the suspension through a 70⍰µm strainer. Following debris removal as per manufacturer’s instructions (Miltenyi Biotec), cells were incubated with CD45^+^ microbeads for 15 min at 4 °C in the dark. CD45^+^ cells were enriched using magnetic-activated cell sorting (MACS) and MS columns. Upon cell counting, samples were fixed using the 10x permeabilization kit and were kept at 4 °C for 16-24 h. Following centrifugation, the supernatant was removed, and cells were resuspended in a solution of glycerol and quenching buffer. All samples were stored at -80 °C until further processing at the Dresden Genome Center (CRTD, Dresden, Germany).

### Processing scRNA-seq data

The reference database for Cell Ranger was built using the mouse reference transcriptome (GRCm39) and gene annotation from Ensembl release 104. The annotation was filtered using *mkgtf* and the genome sequence, along with the filtered annotation, were used as input to *mkref* of Cell Ranger. The raw sequencing data was processed with *count* of Cell Ranger software (v7.1.0) provided by 10x Genomics. Single-cell RNA sequencing (scRNA-seq) data was analyzed using Seurat v5. Cell-type identities assigned through clustering and annotation^59^ and by referencing glioblastoma datasets, including the glioblastoma reference atlas^23,60^. Off-target cells were excluded, and the remaining cells were visualized using UMAP. Specific subsets such as tumor-associated macrophages (TAMs) and T cells were further analyzed. For data normalization and dimensionality reduction, the SCTransform function was applied to 20,000 cells, by retaining all genes. Principal component analysis (PCA) was performed, followed by UMAP visualization in 2D space using the first 30 principal components. Neighbor identification and clustering were conducted using the same PCA dimensions. Differentially expressed marker genes (DEGs) for each cluster were identified using the Wilcoxon rank-sum test and Benjamini-Hochberg method to correct for multiple comparisons (presto v1.0.0). DEGs expressed in at least 25% of cells and showing a log2 fold change > 1 were included.

### Spatial transcriptomics (Visium)

Brain from glioma-bearing mice were harvested, snap-frozen in isopentane and embedded in pre-chilled OCT (Tissue-Tek). Samples were processed under RNAse-free conditions according to the manufacturer’s guidelines (10x protocols, CG000240, CG000160, CG000238, CG00024, and CG000239). 10 µm thick brain sections were obtained from samples with RIN > 7, placed on Visium Spatial Tissue Optimization Slides (PN-3000394) or Visium Spatial Gene Expression slides (PN-2000233) and were sectioned as previously described^61^. Subsequent to imaging and coverslip removal, slides were placed in a Visium Cassette (PN 2000281), and the sections were permeabilized for 18 min. Elution from the capture areas was performed according to manufacturer’s protocols (CG000239) after 0.1X SSC washes, reverse transcription and cDNA second strand synthesis. Upon cDNA amplification (15 - 16 cycles), samples underwent 0.6x purification with SPRIselect beads (Beckman Coulter) to enrich cDNA fragments (> 400 bp) and a quality control on the Fragment Analyzer (using the DNF-473 NGS Fragment Kit, Agilent). The Spatial Gene Expression Library Construction was performed according to the manufacturer’s guidelines. Upon quantification, libraries were sequenced on an Illumina Novaseq6000 in paired-end mode (R1/R2: 100 cycles; I1/I2: 10 cycles), generating from 220 to 270 million fragment pairs per library. Raw sequencing data was processed with *count* of the Space Ranger v2.1.0 software (10x Genomics^62^). To build the reference for Space Ranger, the mouse genome (GRCm39) and gene annotation (Ensembl 104) downloaded from Ensembl were processed following the steps provided by 10x Genomics. Manually, microscope images were aligned to fit the fiducial markers of VisiumHD slides using the Loupe Browser (10x Genomics) and were provided as input for Space Ranger.

Spatial transcriptomics data generated with the 10x Genomics Visium platform were analyzed using Seurat v5^63^ and each tissue slice was processed individually. Raw count matrices were normalized by *SCTransform*, followed by PCA and UMAP visualization using the first 30 principal components. Neighboring spots and clusters were identified by *FindNeighbors and FindClusters* applied to these 30 PCA dimensions. All individually processed slices were merged into a single Seurat object for downstream analysis. Cell-type annotation was performed by integrating unsupervised clustering results with marker gene analysis and label transfer from a published GL261 glioblastoma single-cell reference atlas^22^. To predict the spatial distribution of immune cell populations, annotations for tumor-associated macrophages (TAMs) and T cells were transferred from scRNA-seq datasets of immune cells isolated from glioma tissues of EGFL7+/+ and EGFL7-/- mice to Visium data using *FindTransferAnchors* and *TransferData*. The resulting prediction scores were used to visualize and localize immune cell types across defined spatial domains, including tumor core, tumor shells and brain regions. All analyses were done at the Dresden Genome Center (CRTD, Dresden, Germany).

### Human data

EGFL7 mRNA expression data of glioma specimens were obtained from the Chinese Glioma Genome Atlas (CGGA; http://www.cgga.org.cn) and stratified by IDH mutation status. Expression values (transcripts per million) were log2-transformed. EGFL7 expression and overall survival data for glioblastoma patients were analyzed using the R2 Genomics Analysis and Visualization Platform (https://r2.amc.nl). A correlation between *EGFL7* expression and infiltration of different immune cells was assessed using the tumor immune estimation resource (TIMER; http://timer.canceromics.org/) across glioblastoma cohorts from TCGA pan-cancer datasets. Spearman rank correlations (ρ, p-values) were calculated for bulk RNA sequencing data and adjusted for tumor purity.

### Statistics

All data were analyzed using GraphPad Prism 6 software (GraphPadSoftware, La Jolla, CA, USA) and are shown as mean ± standard error of mean (SEM). Statistical analysis was performed using one-way analysis of variance (ANOVA) for three or more groups followed by Bonferroni hypothesis testing. Mann-Whitney U test was used to analyze significant differences between two groups. For survival analyses, Kaplan-Meier curves were generated and compared using log-rank test.

## Extended Data

**Extended Data Figure 1:**
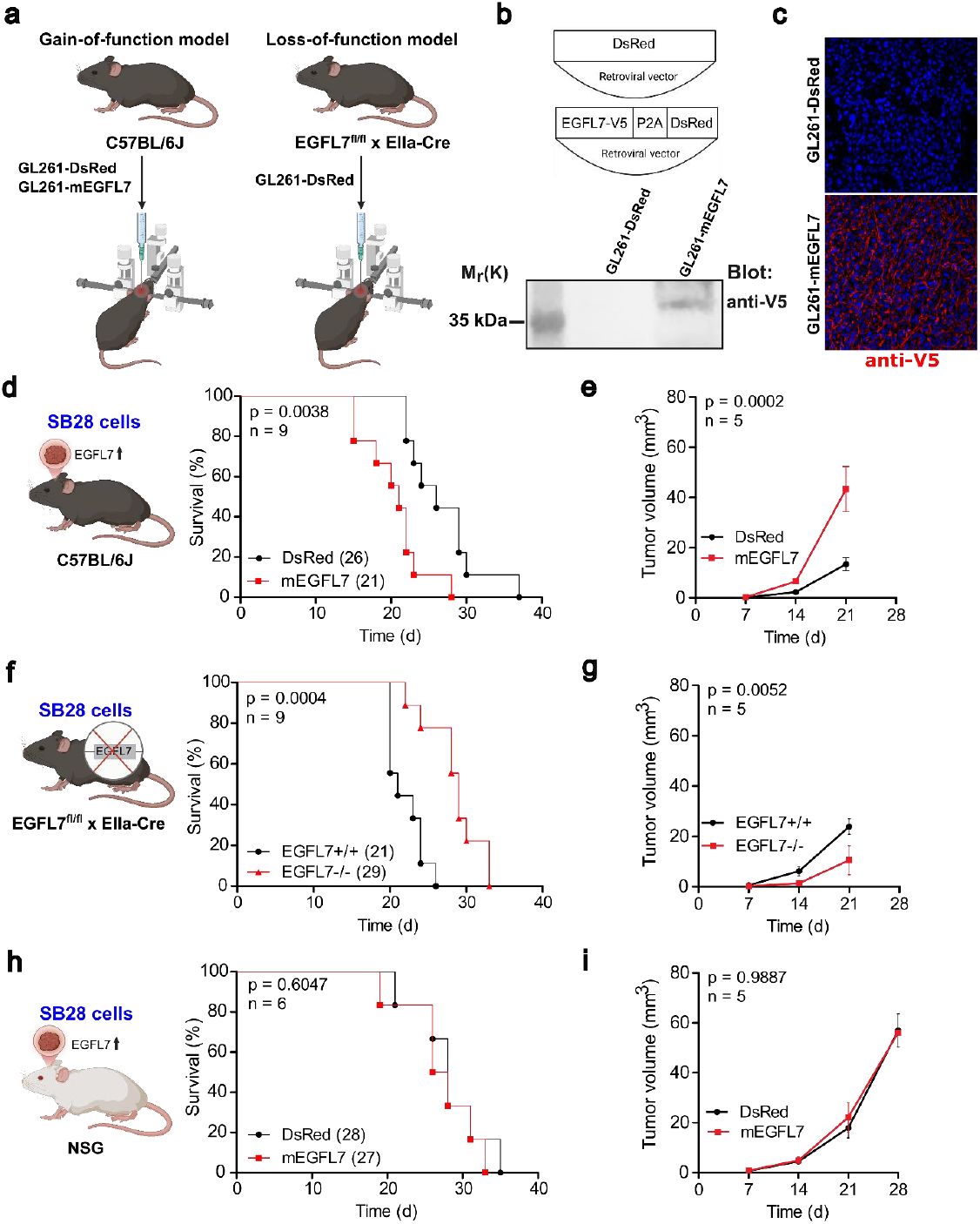
EGFL7 affects survival in glioblastoma. **a:** Schematic of EGFL7 gain- and loss-of-function glioma models applied. b: Schematic of retroviral vectors expressing DsRed or mEGFL7-P2A-DsRed (top). Western blot verification of ectopic V5-mEGFL7 expression (size about 37 kDa) in infected GL261 cells (bottom) **c:** Immunostaining of ectopic V5-mEGFL7 in infected glioma tissue compared to tumors infected with a negative control virus (encoding for DsRed). **d:** Kaplan-Meier curves show reduced survival of mice bearing SB28-mEGFL7 gliomas compared to SB28-DsRed control tumors. **e:** Longitudinal MRI analyses revealed an increased tumor volume in SB28-mEGFL7 glioma-bearing mice compared to SB28-DsRed tumor-bearing animals, 21 d post tumor implantation. **f:** Kaplan-Meier curves show prolonged survival of EGFL7-/- mice implanted with SB28-DsRed gliomas compared to tumor-implanted littermate controls. **g:** Longitudinal MRI analyses showed decreased tumor volumes in EGFL7-/- mice compared to tumor-implanted control littermates, 21 d post tumor implantation. **h:** Immunodeficient NSG mice bearing SB28-mEGFL7 gliomas and SB28-DsRed control tumors survive comparably long. **i:** Longitudinal MRI analyses reveal comparable tumor sizes of SB28-mEGFL7 and SB28-DsRed glioma implanted in NSG mice, 21 d post tumor implantation (right). Statistical analyses were performed using log-rank test (d,⍰f,⍰h) and two-way ANOVA (e,⍰g,⍰i).

**Extended Data Figure 2:**
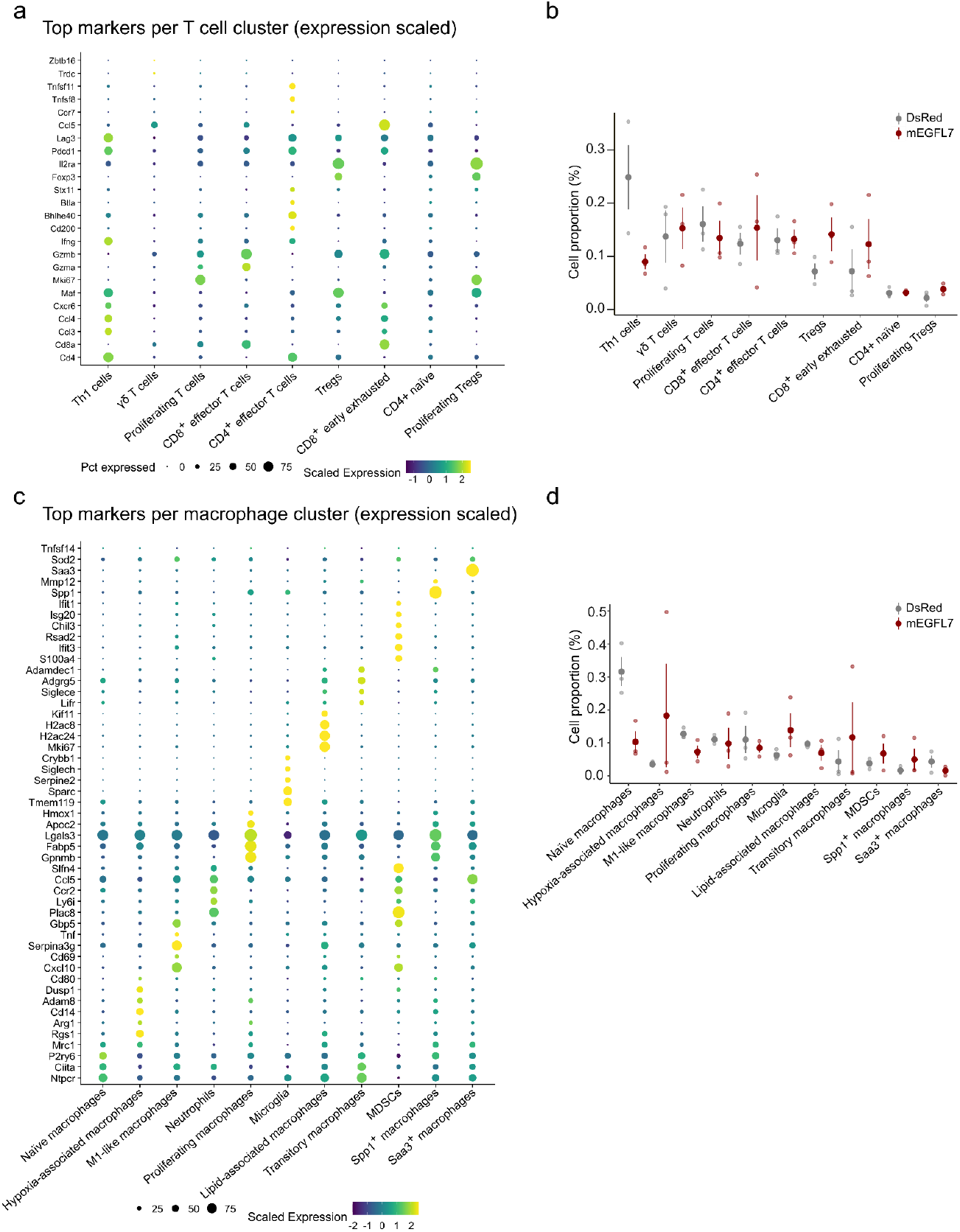
Cell annotation and proportion analysis of ectopic EGFL7’s role in immune cell heterogeneity across biological replicates. **a:** Cell annotation analyses upon scRNA-seq revealed different T cell clusters in GL261-mEGFL7 and GL261-DsRed glioma specimens. **b:** Proportion of T cell subtypes across three biological replicates of GL261-DsRed and GL261-mEGFL7 glioma specimens. **c:** Upon scRNA-seq cell annotation analyses revealed different monocyte/macrophage clusters in GL261-mEGFL7 and GL261-DsRed glioma specimens. d: Proportion of monocyte/macrophage subtypes across 3 biological replicates of GL261-DsRed and GL261-mEGFL7 glioma specimens.

**Extended Data Figure 3:**
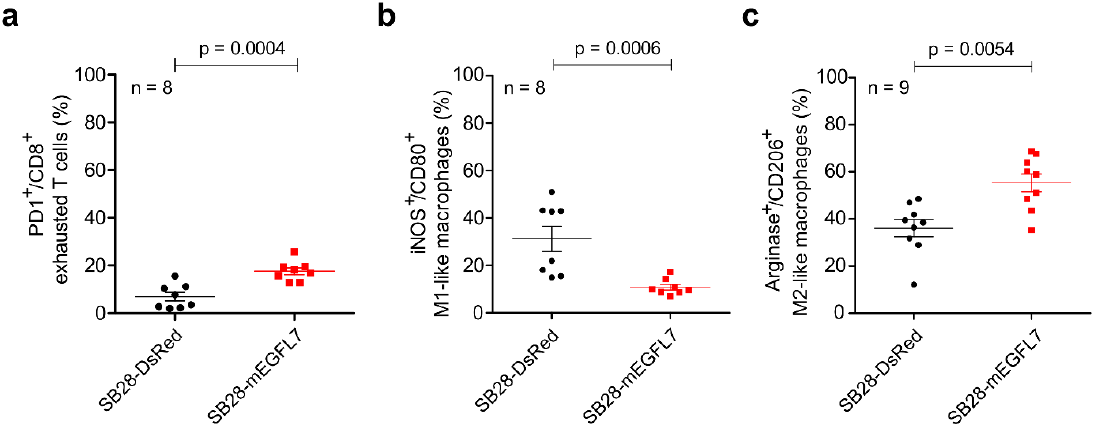
Validation of induced immunosuppression by ectopic EGFL7 expression using SB28 glioma cells. Flow cytometry analyses revealed **a:** an increase in the proportion of CD8^+^ exhausted T cells, **b:** a reduction of M1-like macrophages, and **c:** an increase in M2-like macrophages in SB28-mEGFL7 glioma-implanted mice compared to animals implanted with SB28-DsRed control tumors. Statistical analysis was performed using the Mann-Whitney U test.

**Extended Data Figure 4:**
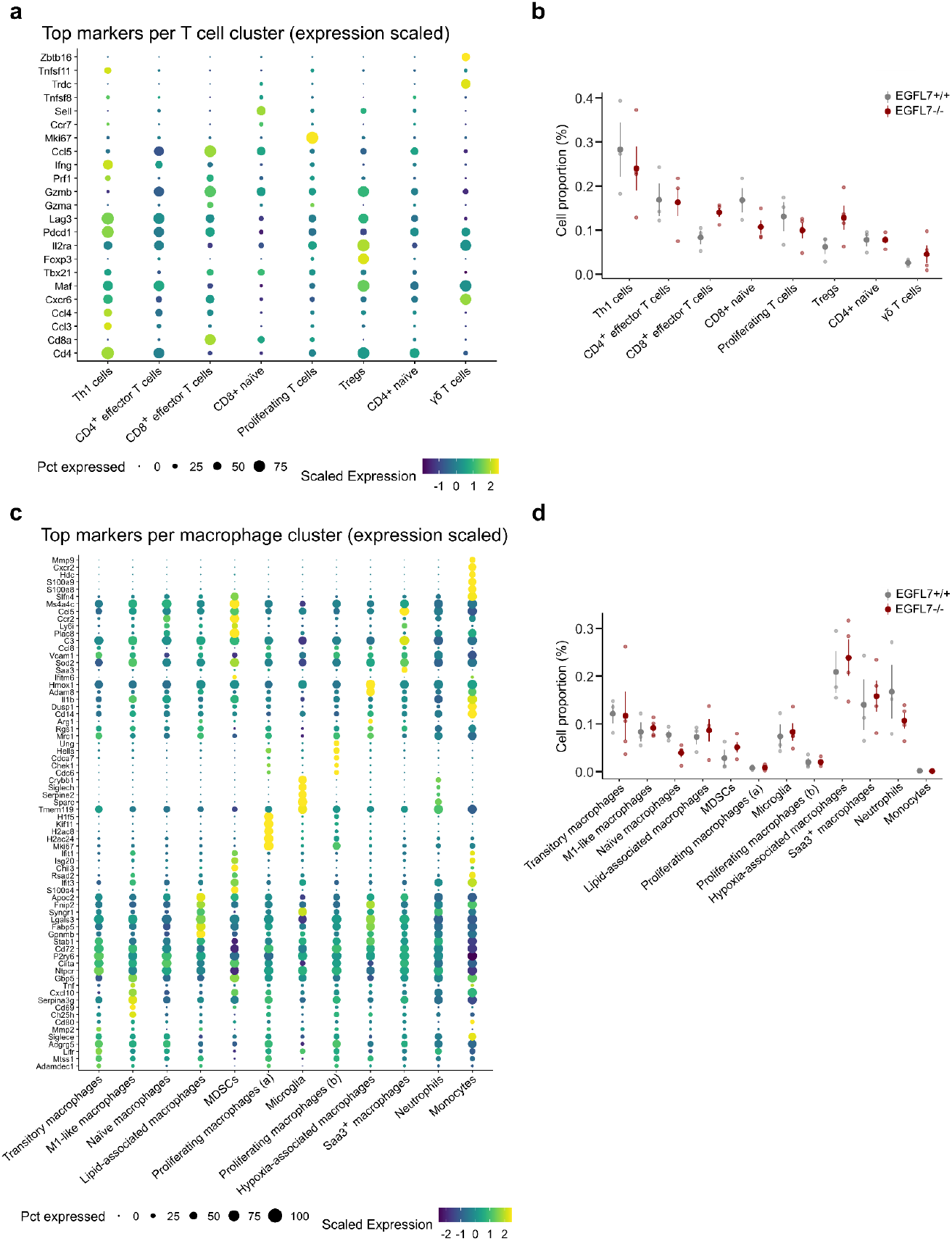
Cell annotation and proportion analysis of immune cells in glioma specimens upon EGFL7 knock-out. **a:** Cell annotation analyses upon scRNA-seq reveal different T cell clusters in GL261-DsRed glioma grown in EGFL7-/- mice and littermate controls. **b:** Proportion of T cell subtypes across three biological replicates per group. **c:** Upon scRNA-seq cell annotation analyses revealed different monocyte/macrophage clusters in GL261-DsRed glioma grown in EGFL7-/- mice and littermate controls. **d:** Proportion of monocyte/macrophage subtypes across 3 biological replicates per group.

## Notes

### Competing Interest Statement

The authors have declared no competing interest.

### Summary of Updates

The revised version includes updated figures, text and author list.

## References

1. van Solinge, T. S., Nieland, L., Chiocca, E. A. & Broekman, M. L. D. Advances in local therapy for glioblastoma — taking the fight to the tumour. Nat. Rev. Neurol. 18, 221–236 (2022).

2. Nørøxe, D. S., Poulsen, H. S. & Lassen, U. Hallmarks of glioblastoma: a systematic review. ESMO Open 1, e000144 (2016).

3. Albini, A. & Sporn, M. B. The tumour microenvironment as a target for chemoprevention. Nature Reviews Cancer vol. 7 (2007).

4. DeCordova, S. et al. Molecular Heterogeneity and Immunosuppressive Microenvironment in Glioblastoma. Frontiers in Immunology vol. 11 (2020).

5. Lee, J., Nicosia, M., Silver, D. & Lathia, J. Sex-specific T cell exhaustion drives differential immune responses in glioblastoma. The Journal of Immunology 210, (2023).

6. Woroniecka, K. et al. T-cell exhaustion signatures vary with tumor type and are severe in glioblastoma. Clinical Cancer Research 24, (2018).

7. Yeo, A. T. et al. Single-cell RNA sequencing reveals evolution of immune landscape during glioblastoma progression. Nat. Immunol. 23, (2022).

8. Mahajan, S., Schmidt, M. H. H. & Schumann, U. The Glioma Immune Landscape: A Double-Edged Sword for Treatment Regimens. Cancers vol. 15 (2023).

9. Hotchkiss, K. M. et al. Dendritic cell vaccine trials in gliomas: Untangling the lines. Neuro-Oncology vol. 25 (2023).

10. Bagley, S. J., Desai, A. S., Linette, G. P., June, C. H. & O’Rourke, D. M. CAR T-cell therapy for glioblastoma: Recent clinical advances and future challenges. Neuro. Oncol. 20, (2018).

11. Aslan, K. et al. Heterogeneity of response to immune checkpoint blockade in hypermutated experimental gliomas. Nat. Commun. 11, (2020).

12. Rizvi, N. A. et al. Mutational landscape determines sensitivity to PD-1 blockade in non-small cell lung cancer. Science (1979). 348, (2015).

13. Hugo, W. et al. Genomic and Transcriptomic Features of Response to Anti-PD-1 Therapy in Metastatic Melanoma. Cell 165, (2016).

14. Ferris, R. L. et al. Nivolumab for Recurrent Squamous-Cell Carcinoma of the Head and Neck. New England Journal of Medicine 375, (2016).

15. Lim, M. et al. Phase III trial of chemoradiotherapy with temozolomide plus nivolumab or placebo for newly diagnosed glioblastoma with methylated MGMT promoter . Neuro. Oncol. 24, 1935–1949 (2022).

16. Ahluwalia, M. S. et al. Phase II trial of SurVaxM combined with standard therapy in patients with newly diagnosed glioblastoma. Journal of Clinical Oncology 36, (2018).

17. Fitch, M.J., et al. Egfl7, a novel epidermal growth factor-domain gene expressed in endothelial cells. Dev. Dyn., 230: 316–324. (2004)

18. Parker, L. H. et al. The endothelial-cell-derived secreted factor Egfl7 regulates vascular tube formation. Nature 428, 754–758 (2004).

19. Dudvarski Stanković, N. et al. EGFL7 enhances surface expression of integrin α 5 β 1 to promote angiogenesis in malignant brain tumors. EMBO Mol. Med. 10, (2018).

20. Larochelle, C. et al. EGFL7 reduces CNS inflammation in mouse. Nat. Commun. 9, (2018).

21. Delfortrie, S. et al. Egfl7 promotes tumor escape from immunity by repressing endothelial cell activation. Cancer Res. 71, 7176–7186 (2011).

22. García-Vicente, L. et al. Single-nucleus RNA sequencing reveals a preclinical model for the most common subtype of glioblastoma. Commun. Biol. 8, 671 (2025).

23. Pombo Antunes, A. R. et al. Single-cell profiling of myeloid cells in glioblastoma across species and disease stage reveals macrophage competition and specialization. Nat. Neurosci. 24, (2021).

24. Li, T. et al. TIMER: A web server for comprehensive analysis of tumor-infiltrating immune cells. Cancer Res. 77, (2017).

25. Xu, H. et al. ITGB2 as a prognostic indicator and a predictive marker for immunotherapy in gliomas. Cancer Immunology, Immunotherapy 71, (2022).

26. Nikolić, I. et al. EGFL7 ligates αvβ3 integrin to enhance vessel formation. Blood 121, (2013).

27. Kim, C. G. et al. VEGF-A drives TOX-dependent T cell exhaustion in anti-PD-1-resistant microsatellite stable colorectal cancers. Sci. Immunol. 4, (2019).

28. Omuro, A. et al. Radiotherapy combined with nivolumab or temozolomide for newly diagnosed glioblastoma with unmethylated MGMT promoter: An international randomized phase III trial. Neuro. Oncol. 25, (2023).

29. Gangoso, E. et al. Glioblastomas acquire myeloid-affiliated transcriptional programs via epigenetic immunoediting to elicit immune evasion. Cell 184, 2454–2470.e26 (2021).

30. Lin, H. et al. Understanding the immunosuppressive microenvironment of glioma: mechanistic insights and clinical perspectives. Journal of Hematology and Oncology vol. 17 (2024).

31. Zhang, H. et al. Roles of tumor-associated macrophages in anti-PD-1/PD-L1 immunotherapy for solid cancers. Molecular Cancer vol. 22 (2023).

32. Reardon, D. A. et al. Effect of Nivolumab vs Bevacizumab in Patients with Recurrent Glioblastoma: The CheckMate 143 Phase 3 Randomized Clinical Trial. JAMA Oncol. 6, (2020).

33. Sampson, J. H. et al. A randomized, phase 3, open-label study of nivolumab versus temozolomide (TMZ) in combination with radiotherapy (RT) in adult patients (pts) with newly diagnosed, O-6-methylguanine DNA methyltransferase (MGMT)-unmethylated glioblastoma (GBM): CheckMate-498. Journal of Clinical Oncology 34, (2016).

34. Bicker, F. et al. Neurovascular EGFL7 regulates adult neurogenesis in the subventricular zone and thereby affects olfactory perception. Nat. Commun. 8, (2017).

35. Barth, K. et al. EGFL7 loss correlates with increased VEGF-D expression, upregulating hippocampal adult neurogenesis and improving spatial learning and memory. Cellular and Molecular Life Sciences 80, (2023).

36. Ravi, V. M. et al. T-cell dysfunction in the glioblastoma microenvironment is mediated by myeloid cells releasing interleukin-10. Nat. Commun. 13, (2022).

37. Azambuja, J. H., Ludwig, N., Yerneni, S. S., Braganhol, E. & Whiteside, T. L. Arginase-1+ exosomes from reprogrammed macrophages promote glioblastoma progression. Int. J. Mol. Sci. 21, (2020).

38. Gérard, A., Cope, A. P., Kemper, C., Alon, R. & Köchl, R. LFA-1 in T cell priming, differentiation, and effector functions. Trends in Immunology vol. 42 (2021).

39. Blythe, E. N., Weaver, L. C., Brown, A. & Dekaban, G. A. β2 Integrin CD11d/CD18: From Expression to an Emerging Role in Staged Leukocyte Migration. Frontiers in Immunology vol. 12 (2021).

40. Yakubenko, V. P. et al. The role of integrin αDβ2 (CD11d/CD18) in monocyte/macrophage migration. Exp. Cell Res. 314, (2008).

41. Haake, M. et al. Tumor-derived GDF-15 blocks LFA-1 dependent T cell recruitment and suppresses responses to anti-PD-1 treatment. Nat. Commun. 14, (2023).

42. Cui, K., Ardell, C. L., Podolnikova, N. P. & Yakubenko, V. P. Distinct migratory properties of M1, M2, and resident macrophages are regulated by αdβ2and αmβ2integrin-mediated adhesion. Front. Immunol. 9, (2018).

43. Pinte, S. et al. Endothelial cell activation is regulated by epidermal growth factor-like domain 7 (Egfl7) during inflammation. Journal of Biological Chemistry 291, 24017–24028 (2016).

44. Helena, J. J. van R. et al. The hippo pathway component taz promotes immune evasion in human cancer through PD-L1. Cancer Res. 78, (2018).

45. Yee, N. K. & Hamerman, J. A. β2 integrins inhibit TLR responses by regulating NF-κB pathway and p38 MAPK activation. Eur. J. Immunol. 43, (2013).

46. Chim, S. M. et al. EGFL7 Is Expressed in Bone Microenvironment and Promotes Angiogenesis via ERK, STAT3, and Integrin Signaling Cascades. J. Cell. Physiol. 230, 82–94 (2015).

47. Xia, T. et al. Advances in the role of STAT3 in macrophage polarization. Frontiers in Immunology vol. 14 (2023).

48. Tharp, K. M. et al. Tumor-associated macrophages restrict CD8+ T cell function through collagen deposition and metabolic reprogramming of the breast cancer microenvironment. Nat. Cancer 5, (2024).

49. Offer, S. et al. Extracellular lipid loading augments hypoxic paracrine signaling and promotes glioma angiogenesis and macrophage infiltration. Journal of Experimental and Clinical Cancer Research 38, (2019).

50. Uhlén, M. et al. A human protein atlas for normal and cancer tissues based on antibody proteomics. Molecular and Cellular Proteomics 4, (2005).

51. García-Carbonero, R. et al. Randomized Phase II Trial of Parsatuzumab (Anti-EGFL7) or Placebo in Combination with FOLFOX and Bevacizumab for First-Line Metastatic Colorectal Cancer. Oncologist 22, (2017).

52. Lakso, M. et al. Efficient in vivo manipulation of mouse genomic sequences at the zygote stage. Proc. Natl. Acad. Sci. U. S. A. 93, (1996).

53. Klaus, T. et al. Impaired Treg-DC interactions contribute to autoimmunity in leukocyte adhesion deficiency type 1. JCI Insight 7, (2022).

54. Göthert, J. R. et al. In vivo fate-tracing studies using the Scl stem cell enhancer: Embryonic hematopoietic stem cells significantly contribute to adult hematopoiesis. Blood 105, (2005).

55. Schmidt, M. H. H. et al. Epidermal growth factor-like domain 7 (EGFL7) modulates Notch signalling and affects neural stem cell renewal. Nat. Cell Biol. 11, 873–880 (2009).

56. Soneoka, Y. et al. A transient three-plasmid expression system for the production of high titer retroviral vectors. Nucleic Acids Research vol. 23 (1995).

57. Demichev, V., Messner, C. B., Vernardis, S. I., Lilley, K. S. & Ralser, M. DIA-NN: neural networks and interference correction enable deep proteome coverage in high throughput. Nat. Methods 17, (2020).

58. Eden, E., Navon, R., Steinfeld, I., Lipson, D. & Yakhini, Z. GOrilla: A tool for discovery and visualization of enriched GO terms in ranked gene lists. BMC Bioinformatics 10, (2009).

59. Abe, P. et al. Molecular programs guiding arealization of descending cortical pathways. Nature 634, (2024).

60. Ruiz-Moreno, C. et al. Charting the single-cell and spatial landscape of IDH-wild-type glioblastoma with GBmap. Neuro. Oncol. 27, (2025).

61. Ståhl, P. L. et al. Visualization and analysis of gene expression in tissue sections by spatial transcriptomics. Science vol. 353 (2016).

62. Zheng, C. et al. Landscape of Infiltrating T Cells in Liver Cancer Revealed by Single-Cell Sequencing. Cell 169, (2017).

63. Hao, Y. et al. Dictionary learning for integrative, multimodal and scalable single-cell analysis. Nat. Biotechnol. 42, (2024).

